# Exploring the Space of Tumor Phylogenies Consistent with Single-Cell Whole-Genome Sequencing Data

**DOI:** 10.64898/2026.01.21.700922

**Authors:** Samin Rahman Khan, Palash Sashittal

## Abstract

Tumors comprise subpopulations of cells that harbor distinct collections of somatic mutations, ranging from single-nucleotide variants (SNVs) to large-scale copy-number aberrations (CNAs). Single-cell whole-genome sequencing (scWGS) enables direct measurement of these mutations; however, inferring tumor phylogenies from scWGS data remains challenging due to ultra-low coverage (∼0.05 ×). There may be multiple ways of imputing missing information in the data leading to distinct tumor phylogenies that are equally well supported by the data. Existing methods produce a single phylogeny and overlook this uncertainty in reconstructing evolutionary histories from sparse scWGS data. We present SCOPE, a novel algorithmic framework that characterizes the space of tumor phylogenies consistent with scWGS data under a copy-number constrained version of the perfect phylogeny model. Our approach relies on estimating the cell fraction of each mutation, i.e. the proportion of cells within each copy-number cluster that carry the mutation. We derive the necessary and sufficient conditions these fractions must satisfy to admit a copy-number constrained perfect phylogeny. This yields a complete combinatorial description of all tumor phylogenies that are supported by the data under our model. We prove that identifying the largest subset of mutations with cell fractions satisfy model constraints using noisy measurements of cell fractions is NP-hard. On simulated data, SCOPE outperforms existing methods in accuracy with faster runtime in particular on the larger simulations. On scWGS data from a patient-derived ovarian cancer cell line, SCOPE infers a more resolved phylogeny with stronger statistical support compared to existing methods. Using SCOPE to analyze a larger dataset of 4 triple negative breast cancer (TNBC) and 8 high-grade serous ovarian cancer (HGSOC) samples, we show that several samples admit multiple phylogenies. We further find that number of admissible phylogenies increases with lower sequencing coverage and is negatively correlated with the number of copy-number clusters and number of distinct loss of heterozygosity (LOH) events in the clusters, highlighting how data quality and evolutionary constraints jointly shape uncertainty in tumor phylogeny reconstruction. By providing a principled framework for exploring and quantifying phylogenetic uncertainty, SCOPE establishes a new foundation for robust inference of tumor evolution from scWGS data.

**Code availability:** Software is available at https://github.com/sashittal-group/SCOPE

## 1. Introduction

Cancer is an evolutionary process in which somatic mutations, such as single-nucleotide variants (SNVs) and copy-number aberrations (CNAs), accumulate in a population of cells. This process results in a heterogeneous tumor with subpopulations of cells, called clones, with distinct genomes. Reconstruction of the evolutionary history of cancer clones, known as a tumor phylogeny, from genomic sequencing data of the cells in a tumor is crucial for understanding cancer progression and developing effective therapies for treatment [1–3].

Single-cell DNA sequencing (scDNA-seq) has transformed our ability to investigate tumor heterogeneity by enabling the measurement of mutations in thousands of individual cells within a tumor. There are two classes of scDNA-seq technologies with different capabilities from measuring SNVs and CNAs. Targeted scDNA-seq technologies [4–7] achieve high sequencing depth (∼50 × per cell) across selected regions of the genome, typically covering cancer-associated genes. This allows accurate identification of SNVs, while detecting CNAs is more challenging. In contrast, single-cell whole-genome sequencing (scWGS) [8–10] provides roughly uniform coverage across the entire genome, enabling reliable detection of CNAs. However, this comes at the cost of ultra low coverage at any particular locus (∼ 0.02–0.1× per cell), which makes detection of SNVs at single-cell resolution challenging.

A comprehensive view of tumor evolution requires characterizing both copy-number aberrations (CNAs) and single-nucleotide variants (SNVs) within a unified phylogenetic framework. Several methods have been developed to infer phylogenies with both SNVs and CNAs from targeted scDNA-seq data [11–14]. However, the reliance of targeted scDNA-seq technologies on pre-defined targets limits their applicability, as novel or unexpected mutations outside the chosen regions remain undetected. In contrast, single-cell whole-genome sequencing provides an unbiased measurement of the tumor mutations without prior assumptions. Moreover, while early analyses of scWGS focused only on inferring CNAs, subsequent studies demonstrated that SNVs could also be detected by pooling information across multiple cells to mitigate sparse coverage [8, 10].

Several methods have been developed to identify SNVs and tumor phylogenies from scWGS data. Clustering-based methods, such as SBMclone [15] and SECEDO [16], identify SNVs and partition cells into clones based on shared mutations. However, these methods do not incorporate the copy-number information nor do they identify a tumor phylogeny on the identified SNV clones. More recently, phylogeny inference-based methods, Phertilizer [17] and Pharming [18], have been developed that simultaneously infer SNV clones and tumor phylogeny under some evolutionary model.

There are two major limitations that are not addressed by the aforementioned methods. First, current methods do not address the ambiguity in the tumor phylogeny inference introduced by the low sequencing coverage of scWGS data. Specifically, there may be multiple ways of imputing missing information in the data leading to distinct tumor phylogenies that are equally well supported by the data. All phylogenies consistent with the scWGS data must be considered to avoid any bias in downstream analyses using the inferred tumor phylogenies, such as identification mutagenic processes that drive cancer progression [19–22]. Second, current methods do not provide a way to filter out sequencing artifacts [23] and SNVs that do not adhere to the constraints of the evolutionary model. Including such mutations in the phylogeny may lead to incorrect reconstructions of the clonal evolution of the tumor.

We introduce SCOPE, an algorithm that characterizes the space of tumor phylogenies that are supported by scWGS data (Fig. 1). Our approach relies on estimating *cell fractions* of each mutation, i.e. the fraction of cells with the mutation in each copy number cluster. We derive the necessary and sufficient conditions that cell fractions must satisfy if they adhere to a copy-number constrained version of the perfect phylogeny model, which provides a complete characterization of all tumor phylogenies that are supported by the scWGS data. We show that finding the largest subset of mutations that satisfy these constrains when estimates of cell fractions may have errors is NP-hard. Leveraging our characterization, SCOPE solves this problem using a mixed-integer linear program (MILP) and enumerates all phylogenies consistent with the observed scWGS data.

**Fig. 1.**
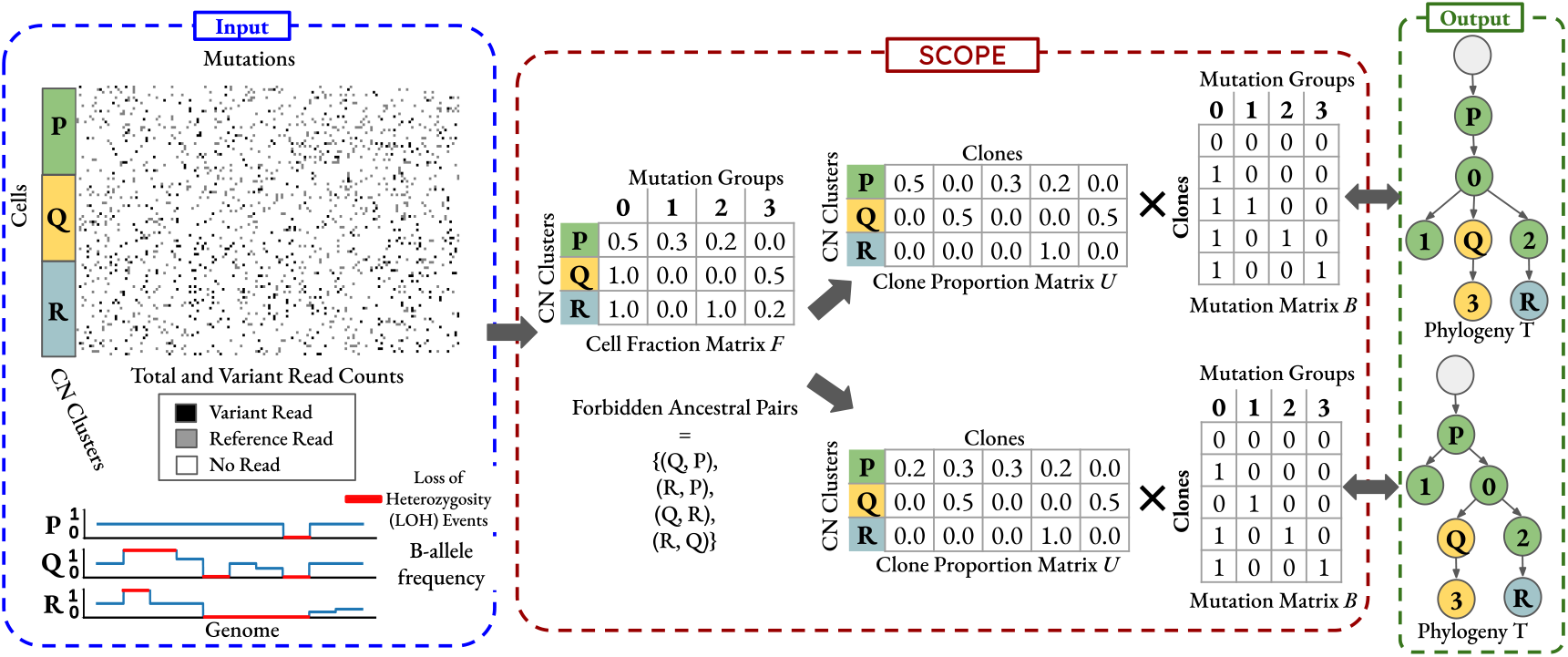
Overview of SCOPE algorithm. (a) Single-cell whole genome sequencing (scWGS) ultra-low coverage measurement of mutations in individual cells which enables reliable measurement of copy-numbers but complicates measurement of SNV clones. SCOPE takes a two-step approach to infer tumor phylogenies from scWGS data. (b) First, SCOPE estimates cell fractions of mutations within each copy-number cluster using a probabilistic read counts model and identifies forbidden ancestral pairs *D* of copy-number clusters. (c) Second, SCOPE enumerates all tumor phylogenies consistent with the estimated cell fractions and forbidden ancestral pairs under a copy-number constrained perfect phylogeny model.

We evaluate SCOPE on both simulated and real datasets to demonstrate its accuracy and its ability to quantify uncertainty in tumor phylogeny inference. On simulated data, SCOPE outperforms existing approaches with faster runtime, especially on large datasets. Applied to scWGS data from ovarian cancer cell lines, SCOPE identifies a single maximal tumor phylogeny consistent with our model constraints, that is more resolved and has stronger statistical support compared to the phylogenies inferred by existing methods. We further analyze a meta-cohort of 4 triple-negative breast cancer and 8 high-grade serous ovarian cancer tumors, showing that uncertainty in phylogeny reconstruction increases as sequencing coverage decreases which reflects the ambiguity introduced by missing data. In contrast, we find that samples with more copy-number clusters and distinct loss-of-heterozygosity (LOH) events in the clusters exhibit fewer admissible solutions, indicating that evolutionary constraints can substantially reduce ambiguity. Together, these results highlight how data quality and evolutionary structure jointly shape uncertainty in scWGS-based phylogeny inference. Finally, we show that SCOPE enables identification of evolutionary relationships that are consistently supported across all admissible phylogenies, in contrast to existing methods that return only a single tree without assessing its robustness.

## 2 Copy-Number Constrained Evolution of Single Nucleotide Variants in Cancer

Cancer is an evolutionary disease that gives rise to a heterogeneous tumor composed of subpopulations of cells called clones carrying distinct sets of mutations. Two major types of genomic alterations drive this evolutionary process: single-nucleotide variants (SNVs), which are point mutations, and copy number aberrations (CNAs), which are amplifications or deletions of contiguous genomic segments. The evolutionary history of the cancer clones is represented by a tumor phylogeny *T*, which is a rooted vertex labeled tree. Each internal vertex *v* of *T* represents an ancestral clone, while the leaves *L*(*T*) represent the extant clones that comprise the tumor. Each vertex is labeled by *b*_*v*_ ∈ {0, 1} ^*m*^ indicating the presence/absence of the *m* mutations. The allele-specific copy-number states of the clones are represented by labeling each vertex with *ĉ*_*v*_ ∈ ℤ^*m*^ and *č*_*v*_ ∈ ℤ^*m*^ indicating the number of copies of each allele of the mutation loci. The root of *T* represents the normal clone that does not have any SNVs and is diploid, i.e. *b*_*r*(*T*),*j*_ = 0, *ĉ*_*r*(*T*),*j*_ = 1 and *č*_*r*(*T*),*j*_ = 1 for all *j ∈* {1, …, *m*}.

Evolutionary models in cancer phylogenetics specify the rules governing how mutations arise in the tumor. The simplest and most widely used of these models is the perfect phylogeny model [24], which assumes that it is highly improbable for the same SNV to occur independently in multiple lineages. Under this model, each mutation arises exactly once and is never subsequently lost. Formally, for each locus *j*, there can be at most one edge (*v, w*) where *b*_*v,j*_ = 0 and *b*_*w,j*_ = 1, and no edge (*v, w*) where *b*_*v,j*_ = 1 and *b*_*w,j*_ = 0. A *perfect phylogeny* is a phylogeny *T* that satisfies these constraints for all mutations.

We extend the perfect phylogeny model by incorporating two types of constraints imposed by the copy-number information of the clones. First, unlike SNVs, identical copy-number states at specific loci can arise independently in different lineages. However, a commonly adopted assumption in models of copy-number evolution is that it is unlikely for entire copy-number profiles to appear more than once independently in the phylogeny [22, 25, 26]. As such, all clones sharing the same copy number profile (*ĉ*_*v*_, *č*_*v*_) must form a connected subtree in the tumor phylogeny. Second, copy-number profiles provide evolutionary constraints through loss-of-heterozygosity (LOH) events in which one allele at a locus is lost due to copy-number deletions, i.e. *ĉ*_*v,j*_ = 0 or *č*_*v,j*_ = 0. Since LOH events are irreversible, i.e. once an allele is lost it cannot be regained, ancestral relationships between certain copy-number clusters becomes biologically infeasible. Specifically, clone *v*′ cannot descend from clone *v* in *T* if *v*′ does not contain all the LOH events present in *v*, i.e. for some *j*, if *ĉ*_*v,j*_ = 0 and *ĉ*_*v*′,*j*_ *>* 0, or if *č*_*v,j*_ = 0 and *č*_*v*′,*j*_ *>* 0.

We introduce these constraints to the perfect phylogeny model and refer to the resulting model as the *copy-number constrained perfect phylogeny model*. Let *p* be the number of distinct clusters of copy-number profiles in the tumor and *forbidden ancestral pairs D* be the set of all pairs (*i, i*′) of copy-number profiles where the set of LOH events in *i*′ is not a subset of the set of LOH events in *i*. We formally define copy-number constrained perfect phylogeny as follows.

### Definition 2.1.

*A copy-number constrained perfect phylogeny T is a tumor phylogeny where*,

1. *Each vertex v is labeled by b*_*v*_ *∈* {0, 1}^*m*^ *and a copy number cluster σ*(*v*) *∈* {1, …, *p*}.
2. *For each mutation j, there is exactly one edge* (*v, w*) *such that b*_*v,j*_ = 0 *and b*_*w,j*_ = 1, *and no edge such that b*_*v,j*_ = 1 *and b*_*w,j*_ = 0.
3. *For any copy-number cluster i, the set of vertices labeled σ*(*v*) = *i form a connected subtree of T* .
4. *For any forbidden ancestral pair* (*i, i*′) ∈ *D of copy-number clusters, there must not be vertices u ≺ u*′ *in T such that σ*(*u*) = *i and σ*(*u*′) = *i*′.

Suppose we have the mutation matrix *B* ∈ {0, 1} ^*r*_×_*m*^ representing the presence/absence of *m* mutations in *r* clones and a copy number clustering *σ* of these *r* clones. We say *T* is a copy-number constrained perfect phylogeny for *B* and *σ* if each leaf of *T* is labeled by a unique row *b*_*ℓ*,:_ of *B* and cluster *σ*(*ℓ*). If a copy-number constrained perfect phylogeny *T* satisfies constraint (4) for a set *D* of forbidden ancestral pairs, then we say *T* is *consistent* with *D*.

## 3 Inferring Tumor Phylogenies from Single-Cell Whole Genome Sequencing Data

Suppose we measure *m* genomic loci in *n* cells using ultra-low coverage (∼0.05 ×) single-cell whole genome sequencing (scWGS). At such low-coverage, it is not possible to reliably determine the presence/absence of SNVs at the measured loci in the cells. For each cell *s* and locus *j*, we observe variant and total read counts; however, most loci will have at most one sequencing read, resulting in extreme sparsity and substantial uncertainty in the mutation status of each cell (Fig. 1a). In contrast, due to the uniformity of the coverage across the genome, it is possible to recover large-scale copy-number profiles by pooling reads across genomic regions. Several methods exist [10, 27–30] to cluster the cells based on their copy number profiles, resulting in a *p × m* allele-specific copy number matrices *C* and *C*, where entries *ĉ*_*i,j*_ and *č*_*i,j*_ denotes the copy number of the two alleles locus *j* in copy number cluster *i*.

Since estimating the presence/absence of mutations in individual cell is highly uncertain at ultra-low coverage, a common practice is to aggregate sequencing reads across cells within each copy-number cluster to form pseudo-bulk samples. Pooling reads in this way increases coverage and enables more reliable estimation of the *cell fraction* of each mutation, i.e. the proportion of cells in a cluster that harbor a given SNV. Several methods [31–35] have been developed to estimate cell fractions from bulk sequencing data, and similar strategies have recently been applied to infer cell fractions from pseudo-bulk scWGS data [8, 18].

We adopt a two-step approach to infer tumor phylogenies from ultra–low coverage scWGS data. First, we estimate cell fractions for each mutation within each copy-number cluster using a probabilistic read-count model (Fig. 1b). We also derive forbidden ancestral copy-number cluster pairs from allele-specific copy-numbers of the clusters. Second, we infer the tumor phylogeny under the copy-number constrained perfect phylogeny model using the estimated cell fractions and forbidden ancestral pairs (Fig. 1c). In the following, we formalize the problem of inferring a tumor phylogeny from estimated cell fractions that is consistent with the forbidden ancestral pairs. We begin with the idealized case where cell-fraction estimates are error-free and then extend the formulation to account for uncertainty in these estimates.

Suppose we have an error-free measurement of the cell fraction matrix *F∈* [0, 1] ^*p*×*m*^, where each entry *f*_*i,j*_ is the fraction of cells in copy number cluster *i* with mutation *j*. The cell fractions are determined by the underlying tumor phylogeny *T* and the proportions of its clones in each copy-number cluster. Let *r* be the number of extant clones in the tumor, i.e. the number of leaves in the tumor phylogeny *T* . We define a *p × r* clone proportion matrix *U*, where each entry *u*_*i,ℓ*_ *∈* [0, 1] is the proportion of cells in copy-number cluster *i* that come from clone *ℓ ∈ L*(*T*), and since these are proportions, we have ∑_*ℓ*_*u*_*i,ℓ*_ = 1. Under the copy-number constrained perfect phylogeny model, the clone proportions must be consistent with the copy-number clustering *σ* of the clones, i.e. we must have *u*_*i,ℓ*_ = 0 if the clone does not belong to the copy-number cluster, i.e. *σ*(*ℓ*) ≠ *i*. This motivates the following problem statement to build a tumor phylogeny from the cell fraction matrix.

### Problem 3.1.

*[Constrained Perfect Phylogeny Mixture (CPPM)] Given cell fraction matrix F and forbidden ancestral pairs D, find phylogeny T, clone proportion matrix U, mutation matrix B and copy number clustering σ such that (i) F* = *UB, (ii) T is a constrained perfect phylogeny for B and σ that is consistent with D and (iii) u*_*i,ℓ*_ = 0 *if σ*(*ℓ*) *≠ i*.

If a cell fraction matrix *F* admits a solution to the CPPM problem with copy-number constrained perfect phylogeny *T*, we say that *T generates F* .

There are two simplifications of the above problem formulation that may not hold in practice. First, due to sequencing errors and sparsity of scWGS data, we do not get error-free estimates of the cell fraction matrix *F* . Second, not all mutations may adhere to the copy-number constrained perfect phylogeny model. Some measured mutations might be the outcome of sequencing artifacts [22, 23], while some true mutations may exhibit violations of the model such as homoplasy (parallel mutations) [36, 37] or losses due to copy-number deletions [11, 13, 38]. To address both the uncertainty in measurement and model violations, we formulate the problem of inferring a copy-number constrained perfect phylogeny with the largest subset of mutations such that the cell fraction *f*_*i,j*_ of each mutation *i* in copy-number cluster *j* is within a given range with lower bound 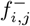 and upper bound 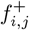. Given lower and upper bound matrices, *F*^−^ and *F* ^+^, and forbidden ancestral pairs *D*, the copy-number constrained perfect phylogeny mixture problem with errors (CPPM-E) problem is posed as follows.

### Problem 3.2.

*[CPPM with Errors (CPPM-E)] Given matrices F*^−^, *F* ^+^ *and forbidden ancestral pairs D, find a largest subset of mutations M*′ ⊆ {1, … *m*} *that admits a p ×* |*M*′| *cell fraction matrix F, tumor phylogeny T, clone proportion matrix U, mutation matrix B and copy number clustering σ such that (i)* 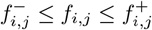 *for each cluster i and mutation j ∈ M*′, *(ii) F* = *UB, (iii) T is a constrained perfect phylogeny for B and σ that is consistent with D and (iv) u*_*i,ℓ*_ = 0 *if σ*(*ℓ*) *?*= *i*.

## 4 Combinatorial Characterization of Tumor Phylogenies Consistent with scWGS Data

We derive a combinatorial characterization of cell fraction matrices that admit a solution to the copy-number constrained perfect phylogeny mixture (CPPM) problem (Prob. 3.1). Specifically, we derive necessary and sufficient conditions that a cell fraction matrix generated by a copy-number constrained perfect phylogeny consistent with a set *D* of forbidden ancestral pairs must satisfy. We also show that the CPPM-E problem (Prob. 3.2) is NP-hard.

Our results build on previous work of characterizing cell fraction matrices generated from perfect phylogenies. This characterization is useful for deconvolving bulk sequencing data to infer tumor phylogenies under the perfect phylogeny model [31, 39–45]. Formally, the perfect phylogeny mixture (PPM) problem involves factorizing the fraction matrix *F* into a clone proportion matrix *U* and a *perfect phylogeny matrix B*, i.e. a mutation matrix that admits a perfect phylogeny.

### Problem 4.1

(Perfect Phylogeny Mixture (PPM)). *Given a cell fraction matrix F, find clone proportion matrix U and perfect phylogeny matrix B such that F* = *UB*.

Cell fraction matrices that admit a solution to the perfect phylogeny mixture problem (Prob. 4.1) can be characterized using a *mutation tree* which describes the order in which mutations arise in the perfect phylogeny. A mutation tree *S* is a rooted tree with *m* + 1 vertices, consisting of the root *v*_0_ and one vertex *v*_*j*_ for each mutation *j* ∈{ 1, …, *m* .} The children of a vertex *v*_*j*_ is denoted by δ_*S*_(*v*_*j*_) and we say mutation *j ≺* _*S*_ *j*′ if *v*_*j*′_ lies on the unique path from the root *v*_0_ to *v*_*j*_ in *S*. El-Kebir et al. [39] showed the a fraction matrix *F* admits a solution to the PPM problem if and only if there exists a mutation tree *S* such that the *sum condition* is satisfied at each vertex, i.e. for each row *i* the cell fraction *f*_*i,j*_ must be greater than the sum of fractions of its children δ_*S*_(*j*) in mutation tree *S*.

### Theorem 4.1

(El-Kebir et al. 2015 [39]). *Cell fraction matrix F* ∈ [0, 1] ^*p*×*m*^ *admits a solution to the PPM problem if and only if it satisfies the sum condition, i*.*e. there exists a mutation tree on the mutations such that* 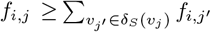 *and* 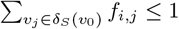 *for all mutations j and rows i*.

Not all frequency matrices that satisfy the sum condition will also admit a copy-number constrained perfect phylogeny mixture problem. This is because the copy-number constrained perfect phylogeny model imposes additional constraints on *U* and *B* using the copy-number information. As such, while the sum condition is necessary, it is not sufficient for a cell fraction matrix to admit a solution to the CPPM problem.

To obtain a complete characterization of frequency matrices that admit a solution to CPPM problem, we derive additional constraints that must be satisfied by *F* with respect to a mutation tree *S*. For each copy-number cluster *i*, a mutation *j* can be classified as clonal (*f*_*i,j*_ = 1), subclonal (0 *< f*_*i,j*_ *<* 1) or absent (*f*_*i,j*_ = 0). We make four key observations. First, since each mutation is gained on exactly one edge in the tumor phylogeny, every mutation can be subclonal in at most one copy-number cluster. Second, for every copy-number cluster, subclonal mutations must be a descendant of clonal mutations in the mutation tree *S*. Third, since each copy-number cluster must form a subtree in the tumor phylogeny, there must be a clone in mutation tree *S* that has a mutation if and only if it is clonal in the copy-number cluster. Fourth, for every forbidden ancestral pair (*i, i*′), a mutation *j* clonal in cluster *i*′ must not be subclonal in cluster *i*. These constraints together with the sum condition yield a complete combinatorial characterization of the frequency matrices that admit a solution to the CPPM problem.

### Theorem 4.2.

*A p × m cell fraction matrix F and forbidden ancestral pairs D admit a solution to the copy-number constrained perfect phylogeny mixture problem if and only if there exists a mutation tree S on the m mutations such that*

1. *for every mutation j, there is at most one row i such that* 0 *< f*_*i,j*_ *<* 1,
2. *for every pair* (*j, j*′) *of mutations, j ≺*_*S*_ *j*′ *if for some row i we have f*_*i,j*_ = 1 *and* 0 *< f*_*i,j*′_ *<* 1,
3. *for every row i, there exists a vertex v in S such that mutation j is on the path from root to v in S if and only if j is clonal in cluster i (f*_*i,j*_ = 1),
4. *for every forbidden ancestral pair* (*i, i*′) ∈ *D, a subclonal mutation j in cluster i (*0 *< f*_*i,j*_ *<* 1*) must not be clonal in cluster i*′ *(f*_*i*′,*j*_ = 1),
5. *F satisfies the sum condition with S*.

A corollary to Condition (3) is that clones defined by the clonal mutations of the copy-number clusters must satisfy the perfect phylogeny constraints. We represent these clones with *binarization F*′ of the cell fraction matrix *F*, which is a binary matrix where entry 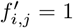 if and only if *j* is clonal in copy-number cluster *i* (*f*_*i,j*_ = 1). We formally state this corollary as follows.

### Corollary 4.1.

*If a copy-number constrained perfect phylogeny T generates a cell fraction matrix F, then its binarization F*′ *is a perfect phylogeny matrix*.

Lastly, we show that the CPPM-E problem in NP-hard. Proofs for theorems and corollaries in this section are provided in Supp. Sec. A.

### Theorem 4.3.

*The CPPM-E Problem is NP-hard*.

## 5 SCOPE: Algorithm to Infer Tumor Phylogenies Consistent with scWGS Data

We develop a novel algorithm, SCOPE (Single-Cell Optimization for Phylogeny Estimation), that uses the characterization derived in the previous section to enumerate all tumor phylogenies consistent with single-cell whole genome sequencing (scWGS) data. SCOPE operates in two stages. First, it estimates confidence interval bounds, *F*^−^ and *F* ^+^, for cell fractions of mutations using a probabilistic model (Sec. 5.1). Second, it construct all copy-number constrained perfect phylogenies that are solutions to the CPPM-E problem given the estimated *F*^−^ and *F* ^+^ matrices and forbidden ancestral pairs *D* (Sec. 5.2).

### 5.1 Estimating cell fraction from read count data

In this section, we describe a probabilistic framework to estimate cell fractions, i.e. the proportion of cells harboring each mutation, from pseudo-bulk samples derived from copy-number clusters in single-cell WGS data. After pooling reads across cells for each copy-number cluster, we obtain a variant read count matrix *Q* and total read count matrix *R*, where entries *q*_*i,j*_ and *r*_*i,j*_ are the variant and total read counts of locus *j* in cluster *i*, respectively. We provide a general description of the model here, with details in Supp. Sec. B.1.

In order to estimate cell fraction from read counts, we require knowledge of mutation multiplicities, i.e. the number of copies of the mutated allele in the cells. We make two assumptions that are consistent with the copy-number constrained perfect phylogeny model and prior work [11, 22, 35]. First, when a mutation arises, it occurs on exactly one copy of the locus. Second, mutation multiplicities can change only through copy-number aberrations. Consequently, all cells in a copy-number cluster *i* that harbor a mutation *j* will have the same multiplicity *x*_*i,j*_, such that 1 ≤ *x*_*i,j*_ ≤ *c*_*i,j*_, where *c*_*i,j*_ is the total copy-number *ĉ*_*i,j*_ + *č*_*i,j*_ of locus *j* in that cluster. Moreover, if mutation *j* is subclonal (0 *< f*_*i,j*_ *<* 1) in cluster *i*, its multiplicity must be *x*_*i,j*_ = 1 and conversely if *x*_*i,j*_ *>* 1, then *j* must be clonal (*f*_*i,j*_ = 1) in cluster *i*.

We model the number *q*_*i,j*_ of variant reads as a binomal random variable: *q*_*i,j*_ ∼Binom(*p*_*i,j*_, *r*_*i,j*_), where *p*_*i,j*_ is the probability of observing a variant read given by *p*_*i,j*_ = *f*_*i,j*_*x*_*i,j*_*/c*_*i,j*_, where *f*_*i,j*_ is the cell fraction of the mutation. For each mutation, we compute the maximum likelihood estimate 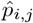 under this Binomial model and use it to infer the cell fractions. Intuitively, since *x*_*i,j*_ = 1 and *f*_*i,j*_ *<* 1 for subclonal mutations, we can identify a threshold *θ*_*i,j*_ = 1*/c*_*i,j*_ which bounds *p*_*i,j*_ from above, i.e. *p*_*i,j*_ *< θ*_*i,j*_. As such, if we observe 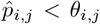, we identify the mutation to be subclonal and set 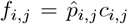, and when 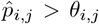, we identify the mutation to be clonal and set *f*_*i,j*_ = 1. We quantify the uncertainty in our estimates by clustering mutations into groups based on the estimated cell fractions and computing empirical lower 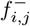 and upper 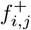 quantiles of the distribution of cell fractions for each mutation group across the cells in each copy-number cluster (details in Supp. Sec. B.1).

### 5.2 Mixed integer linear program for phylogeny estimation

We formulate a mixed integer linear program (MILP) that uses the characterization derived in Section 4 to generate all solutions to the CPPM-E for given bounds *F*^−^ and *F* ^+^ for the cell fraction matrix *F* and a set *D* of forbidden ancestral copy-number cluster pairs. Specifically, the MILP infers *F* that satisfies the bounds (*F*^−^ ≤ *F* ≤ *F* ^+^) and the constraints derived in Theorem 4.2. We employ an iterative procedure that progressively adds constraints to the MILP to enumerate multiple solutions to the problem. We also propose a metric to prioritize the multiple solution based on the fit of the inferred cell fraction matrix *F* (details in Supp. Sec. B.3)

## 6 Results

### 6.1 Simulated Data

We compared the performance of SCOPE against SBMClone [15] and Phertilizer [17] on simulated data. We simulated cancer phylogenies with *n* = 1000, 5000, 10000 cells, *p* = 5, 10, 15 copy-number clusters and number *m* of mutations varying from 500 to 15000 using a previously published simulator [11]. The read count data was simulated using a beta-binomial model with uniform coverage of cov = 0.02 ×, 0.05×, 0.1 × and a sequencing error rate *ϵ* = 0.001 (similar to error rates for Illumina sequencing [46]). We simulate 5 instances for each combination of the varying simulation parameters. Detailed description of the simulations are provided in the Supp. Sec. C. We had to exclude Pharming [18] from this simulation study because it was taking more than 30 minutes to infer a phylogeny on each simulation instance.

We evaluate the accuracy of the inferred phylogeny 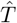 against the ground-truth phylogeny *T* by computing the widely used metric *mutation relation accuracy* 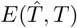 [11, 13, 17, 38]. In a perfect phylogeny, two mutations *j, j*′ may be in exactly one of four ways: (i) *j* is ancestral to *j*′, (ii) *j*′ is ancestral to *j*, (iii) *j* and *j*′ occur on the same branch (iii) *j* and *j*′ occur on distinct lineages in the phylogeny (i.e. they are incomparable). Mutation relation accuracy 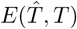measures the fraction of mutation pairs for which these relationships are correctly recovered from the inferred phylogeny 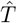. We also evaluate *mutation placement accuracy* 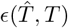, which quantifies the accuracy of assigning mutations as clonal or subclonal within each copy-number cluster. SBMClone does not infer a tumor phylogeny and instead yields a clustering of SNVs and cells with density of variant reads of SNV clusters observed in the cell clusters. Following previous work [15, 22], we derive a perfect phylogeny from SBMClone results by treating an SNV cluster as present in a cell cluster if the density exceeds a specified threshold. Further details of the metrics and how they were computed for each method are provided in the Supplement C.1.

SCOPE outperforms existing methods both in terms of mutation relation accuracy (Fig. 2a) and mutation placement accuracy (Fig. 2b) across all simulation parameters. For the largest simulations with *n* = 10000 cells, SCOPE achieves the highest mutation relation accuracy 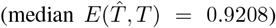 and mutation placement accuracy 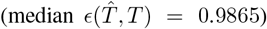 compared to Phertilizer 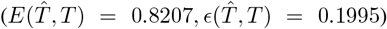, and SBMClone 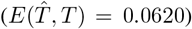. Notably, the superior performance of SCOPE is accompanied with faster runtime compared to Phertilizer on simulations with more than 1000 cells (median 746s vs 5439s for Phertilizer when *n* = 10000, Fig. 2c). This computational efficiency arises from the ability of SCOPE to generate the phylogeny from a *p × m* cell fraction matrix, where the number *p* of copy-number clusters is much smaller than the number *n* of cells. Finally, when SCOPE is provided with the ground-truth mutation groups, i.e. sets of mutations that occur on the same branch in the ground-truth phylogeny, it achieves near perfect accuracy (median 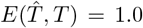, 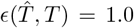, Fig. S3), indicating that improved mutation clustering can further enhance performance of SCOPE.

**Fig. 2.**
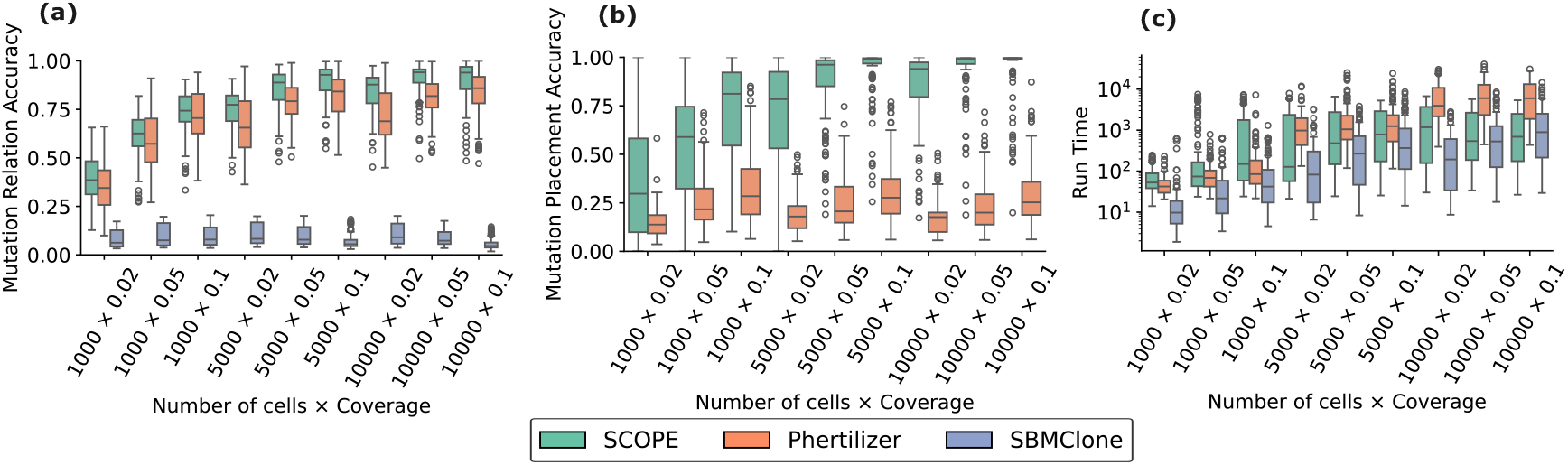
SCOPE outperforms and runs faster than existing methods in recovering the tumor phylogenies on simulated data. (a) Pairwise relationship accuracy, (b) mutation placement accuracy compared to the simulated ground truth, and (c) run-time for each method on simulated data each method across simulated datasets varying in cell number and coverage. Box plots show the median and the interquartile range (IQR), and the whiskers denote the lowest and highest values within 1.5 times the IQR from the first and third quartiles, respectively.

### 6.2 SCOPE infers ovarian cancer phylogeny with strong statistical support

We used SCOPE to scWGS data of *n* = 890 cells from three clonally-related cancer cell lines sourced from the same high-grade serous ovarian cancer (HGSOC) patient [8] with relative high coverage of 0.25 × . Laks et al. [8] identified *m* = 14608 SNVs and *p* = 9 copy-number clusters in the data. SCOPE reveals that there is only one tumor phylogeny with the largest set of mutations (13,379) that satisfy the constrained perfect phylogeny model. The SCOPE phylogeny with the lowest cell fraction distortion has 12 clones (Fig. 3a). We compare the SCOPE phylogeny with phylogenies inferred by Phertilizer, which has 10 clones with 13,832 mutations (Fig. 3b), and the phylogeny reported by Laks *et. al*. in the original publication which has 17 clones with 14,608 mutations (Fig. 3c). We were not able to run Pharming on this data due to the size of the dataset. Mutations were annotated using OncoKB [47] and filtered based on frequencies observed in ovarian cancer datasets on cBioPortal [48] (details in Supp. Sec. F).

**Fig. 3.**
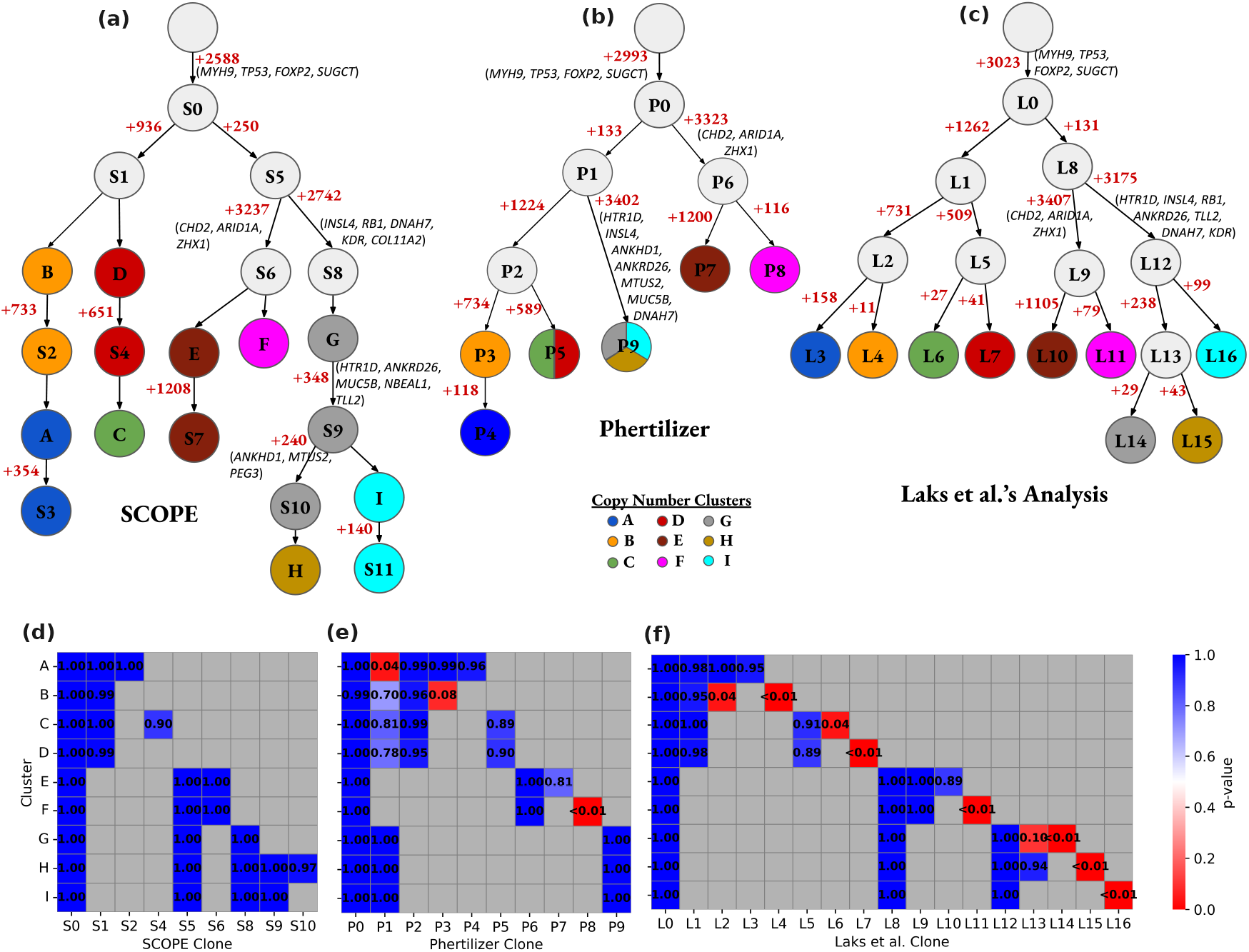
SCOPE infers a more resolved ovarian cancer phylogeny with stronger statistical support than existing methods. Phylogenies inferred by (a) SCOPE, (b) Phertilizer and (c) Laks *et al*. analysis. Statistical support of clonal mutations in the phylogeny inferred by (d) SCOPE, (e) Phertilizer and (f) Laks *et al*. analysis.

SCOPE provides a more resolved view of tumor evolution compared to existing methods by distinguishing between clonal and subclonal mutations in copy-number clusters. SCOPE, Phertilizer and Laks *et al*. analysis agree on the major evolutionary patterns: the same set of driver mutations (*TP53, SUGCT*, and *MYH9*) appear on the trunk, and the phylogenies consistently recover three large clades composed of (i) copy-number clusters A, B, C, D, (ii) clusters E, F, and (iii) clusters G, H, I (Fig. S2). However, unlike the other two methods, SCOPE distinguishes subclonal from clonal mutations within the copy-number clusters. For example, Phertilizer and Laks *et al*. place *ANKRD26, TLL2, RB1, DNAH7, HDR*, and *COL11A2* mutations on the same branch of the (G, H, I) clade, leaving their ordering unresolved. SCOPE refines this region of the tree, inferring that *ANKRD26, MUC5B, NBEAL1, TLL2* occur after *RB1, DNAH7, KDR*, and *COL11A2* in cells from copy-number cluster G, thereby revealing the subclonal structure missed by other methods.

We construct a hypothesis test to evaluate the statistical support for clonal mutations of copy-number clusters in the inferred phylogenies. For a mutation *j* is clonal is copy-number cluster *i*, its expected cell fraction *f*_*i,j*_ = 1. As such, we can model the distribution of variant read counts clonal mutation as a Beta-Binomial with success of probability *p*_*i,j*_ ≥ 1*/c*_*i,j*_, where *c*_*i,j*_ is the total copy-number of locus *j* in cells of copy-number cluster *i*, and δ is the dispersion parameter. Since mutations within a cluster may arise from loci with different copy-number states, we partition each mutation cluster *M* into *M*_1_, …, *M*_*d*_ where each part *M*_*s*_ is the set of mutations *j* such that *c*_*i,j*_ = *s* and *d* is the largest copy-number. The null hypothesis *H*_0_ states that mutation cluster *M* is clonal in copy-number cluster *i*, i.e. *fi,M*_*s*_ = 1 for all *s* = 1, …, *d*. We therefore perform a conjunction of tests 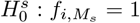 using the Beta–Binomial cumulative mass function to compute individual *p*-values, and combine them using Stouffer’s method [49] (details in Supp. Sec. E).

SCOPE phylogeny shows stronger statistical support of clonal mutations compared to phylogenies inferred by Phertilizer and Laks *et. al*. phylogenies. We applied the same clonality hypothesis test independently to every clonal mutation group identified by each method, resulting in a comparable number of tests across approaches (28 for SCOPE, 32 for Phertilizer, and 37 for Laks *et al*.; Fig. 3d–f). The SCOPE phylogeny shows strong statistical support for clonal mutation clusters. Specifically, none of the 28 null hypotheses for clonality in the SCOPE phylogeny were rejected at a false discovery rate (FDR) of 0.05 (Fig. 3d) indicating strong support for all clonal mutation groups. In contrast, Phertilizer failed 2 of 32 tests (Fig. 3e), with clonality rejected for mutation group P1 in copy-number cluster A (*p*-value= 0.042) and P8 in cluster F (*p* = 1.1 × 10^−54^). The phylogeny from Laks et al. showed even weaker support (Fig. 3f), with 8 of 37 tests rejected, including mutation groups L2 and L4 in cluster B (*p* = 0.042 and 0.003), L6 in cluster C (*p* = 0.042), L7 in cluster D (*p* = 1.3 × 10^−5^), L11 in cluster F (*p* = 4.1 × 10^−7^), L14 in cluster G (*p* = 3.0 × 10^−11^), L15 in cluster H (*p* = 1.1 × 10^−24^), and L16 in cluster I (*p* = 3.3 × 10^−9^). These results suggest that SCOPE provides a more statistically robust reconstruction of tumor clonal structure from scWGS data.

### 6.3 SCOPE quantifies uncertainty in phylogeny inference across a meta-cohort of 12 tumors

We applied SCOPE to a meta-cohort of 12 tumors [19] to investigate the sources of uncertainty in phylogenetic inference. The cohort comprises 4 triple-negative breast cancers (TNBCs) and 8 high-grade serous ovarian cancers (HGSOCs), which are further grouped into three categories based on copy-number mutational signatures: HRD-Dup (enriched in small tandem duplications and BRCA1 mutations), FBI (enriched in fold-back inversions and CCNE1 amplification), and TD (large tandem duplications associated with CDK12 mutations) [19]. Specifically, the cohort includes 4 HGSOC HRD-Dup, 3 HGSOC FBI, 1 HGSOC TD, 2 TNBC HRD-Dup, and 2 TNBC FBI samples. Across

SCOPE reveals how data quality and evolutionary constraints jointly shape the uncertainty in phylogeny inference. As expected, we find the the uncertainty in phylogeny inference, quantified by the number of distinct admissible tumor phylogenies supported by the data, is strongly influenced by sequencing coverage. We observe a negative correlation between coverage and uncertainty in phylogeny inference (Fig. 4a, *R*^2^ = 0.321, Wald Test *p*-value = 0.0549), reflecting the increased ambiguity introduced by missing data. SCOPE also reveals that copy-number heterogeneity of the tumor constrains the solution space and reduces uncertainty under the copy-number constrained perfect phylogeny model. Specifically, uncertainty in phylogeny inference is negatively correlated with number *p* of copy-numbers clusters (Fig. 4b, *R*^2^ = 0.117, *p*-value= 0.276) and, for samples with coverage is less than 0.09 ×, with normalized number |*D*|/*p*^2^ of forbidden ancestral copy-number cluster pairs (Fig. 4c, *R*^2^ = 0.220, *p*-value= 0.203). Together, these results highlight that both the quality of scWGS measurements and the richness of underlying evolutionary constraints determine how much of the phylogeny is uniquely recoverable from the data.

**Fig. 4.**
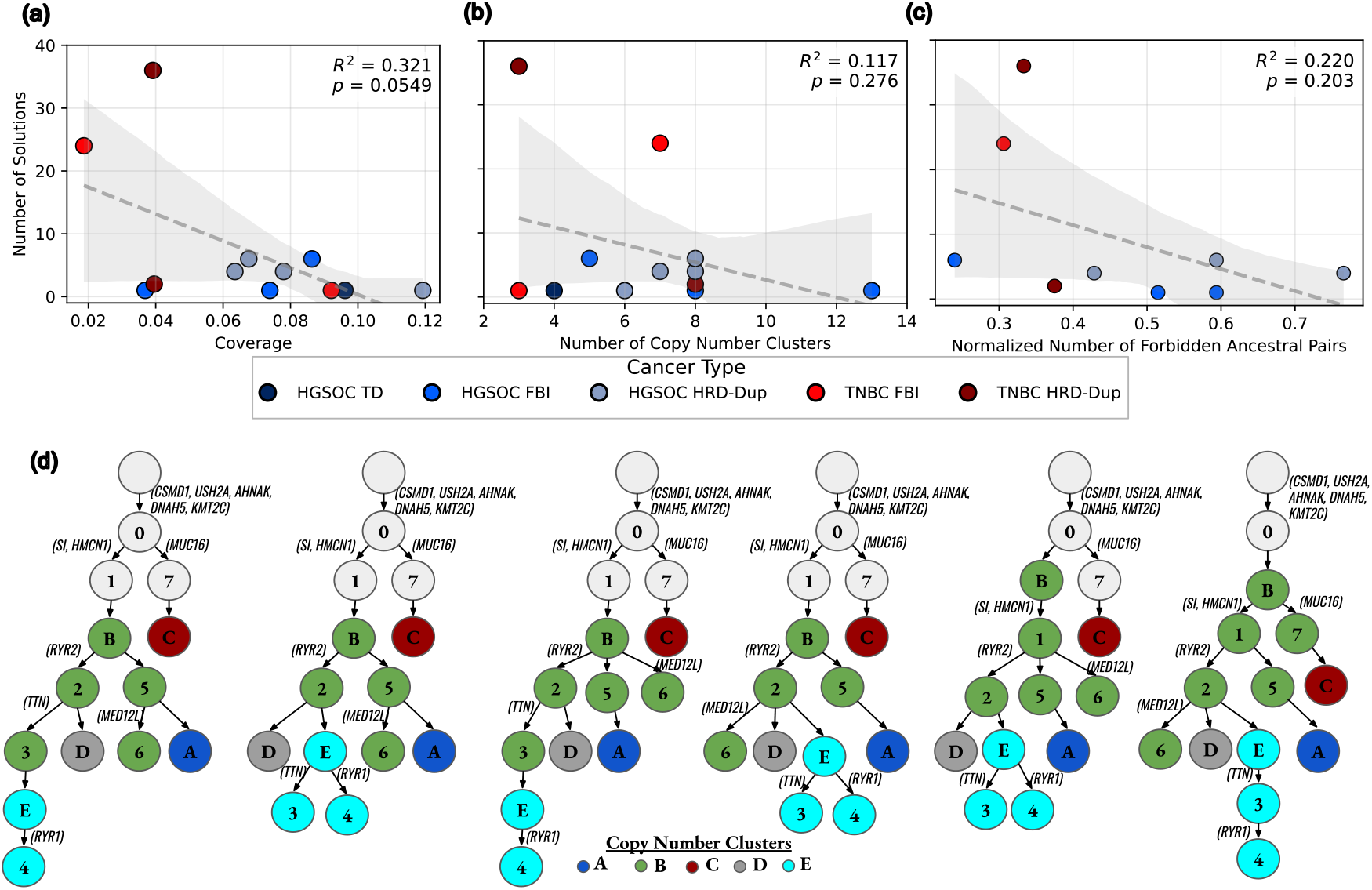
SCOPE uncovers unambiguous evolutionary relationships and reveals sources of uncertainty in phylogeny inference across a meta-cohort of 12 tumors. Number of phylogenies inferred by SCOPE for each tumor across different (a) sequencing coverage, (b) number of copy-number clusters. (c) Number of inferred phylogenies for samples with coverage *<* 0.09 × for varying normalized number |*D*|/*p*^2^ of forbidden ancestral pairs. (d) Six plausible tumor phylogenies for a HGSOC FBI sample (SA1091) inferred by SCOPE. these tumors, SCOPE identifies multiple admissible phylogenies for several samples (median = 3, mean = 7.25). These samples vary widely in sequencing coverage (0.01 × - 0.1×), number of cells (250 – 6, 000), number of mutations (6, 000 – 44, 000), and number of copy-number clusters (3 – 13) (summary of the cohort in Supp. Table S1).

SCOPE enables identification of evolutionary relationships that are unambiguously supported across all admissible phylogenies. We illustrate this using sample SA1091, a HGSOC FBI tumor for which SCOPE identifies six admissible phylogenies involving 7 mutation groups (with 8, 928 SNVs) and 5 copy-number clusters. SCOPE reveals uncertainty in the placement of mutation groups 3 (containing *TTN* mutation), 4 (*RYR1*) and 6 (*MED12L*), and clonality of groups 3 in cluster E and groups 1 (*SI, HMCN1*) and 7 in cluster B. In contrast, mutation group 0, comprising of *CSMD3, USH2A, AHNAK, DNAH5* and *KMT2C* mutations, consistently appears on the trunk, and the relative ordering of groups 0, 1, 7, 2 (*RYR2*) and 5 is preserved across all admissible phylogenies. These results demonstrate that unlike existing methods which only return a single phylogeny without quantifying the uncertainty in the result, SCOPE enables us to distinguish robust evolutionary events from uncertain ones.

## 7 Discussion

We introduce SCOPE, a novel framework to quantify uncertainty in phylogeny inference from ultra-low coverage single-cell whole genome sequencing (scWGS) data. SCOPE relies on a complete characterization of mutation cell fractions, i.e. the fraction of cells within each copy-number cluster that carry a mutation, that are generated by a copy-number constrained perfect phylogeny. Leveraging this characterization, SCOPE infers all tumor phylogenies that are supported by the data. This enables identification of high-confidence evolutionary trajectories that are observed in every plausible phylogeny that explains the data. SCOPE also ranks phylogenies based on the deviation of inferred cell fractions from those observed in the data, helping to identify the most reliable solutions from potentially large solution spaces under high-uncertainty conditions.

We demonstrate the superior performance of SCOPE compared to existing methods on both simulations and real data. On an ovarian cancer dataset [8], SCOPE reconstructs a more resolved and statistically better supported phylogeny than existing methods, revealing subclonal structure that was missed in earlier analyses. On a broader cohort of 4 triple-negative breast cancer and 8 high-grade serous ovarian cancer tumors [19], SCOPE shows that several samples support multiple distinct tumor phylogenies. Notably, we find that the number of admissible phylogenies decreases as sequencing coverage increases, and as the number of copy-number clusters and unique loss of heterozygosity events across these clusters increases. These results highlight the interplay between data quality and evolutionary constraints as key determinants of uncertainty in inferred phylogenies.

This work opens several avenues of future research. First, the evolutionary model underlying SCOPE can be extended to include loss of SNVs due to copy-number deletions and integrated with models of copy-number evolution. Second, SCOPE currently enumerates all possible solutions of the constrained perfect phylogeny mixture problem. Future versions of SCOPE could focus on identifying a consensus phylogenies that capture the features conversed across all solutions without exhaustively enumerating all solutions. Third, SCOPE can be used to systematically investigate how uncertainty in the inferred phylogenies depends on experimental parameters such as coverage and number of sequenced cells. Such analyses could guide future single-cell sequencing experiments by quantifying the trade-off between sequencing cost and uncertainty in the solution space. Lastly, while we applied SCOPE to cancer, the underlying approach of pooling information across cells to overcome sparsity of whole-genome sequencing and quantifying the uncertainty in phylogeny inference is broadly applicable. This framework can be applied in other biological contexts, such as developmental systems, and with other types of sparse sequencing assays such as single-cell RNA sequencing [50] or single-cell ATAC sequencing [51]. We envision that SCOPE will lay the foundation for future development of methods for robust and uncertainty-aware phylogenetic inference from single-cell sequencing data.

