## Supplemental Materials for "Exploring the Space of Tumor Phylogenies Consistent with Single-Cell Whole-Genome Sequencing Data"

**Supplementary**

**D Mutation clustering and forbidden ancestral pairs in real data** 10

**E Statistical test for evaluating clonal relationships** 11

**F Annotating Mutated Genes** 11

**G Method parameters** 12

**H Supplementary results** 12

### A Proofs

#### A.1 Proof to Theorem 4.2 and Collolary 4.1

We restate the theorem here and then provide a proof.

**Theorem A.1.** *A $p \times m$ cell fraction matrix $F$ admits a solution to the copy-number constrained perfect phylogeny mixture problem if and only if there exists a mutation tree $S$ on the $m$ mutations such that*

1. *for every mutation $j$, there is at most one row $i$ such that $0 < f_{i,j} < 1$,*
2. *for every pair $(j, j')$ of mutations, $j \prec_S j'$ if for some row $i$ we have $f_{i,j} = 1$ and $0 < f_{i,j'} < 1$,*
3. *for every row $i$, there exists a vertex $v$ in $S$ such that mutation $j$ is on the path from root to $v$ in $S$ if and only if $f_{i,j} = 1$,*
4. *$F$ satisfies the sum condition with $S$.*

*Proof.* ($\Rightarrow$) We will prove that if a cell fraction matrix $F$ generated by a constrained perfect phylogeny $T$ consistent with forbidden ancestral pairs $D$, then $F$ satisfies the five condition stated in the theorem above. We obtain mutation tree $S$ from phylogeny $T$ as follows. We first collapse all the edges $(u, v)$ where $b_{u,j} = b_{v,j}$ for all $j$. Each vertex is then labeled by the mutation $j$ gained on its incoming edge.

**Condition (1)** Since $T$ is a constrained perfect phylogeny, each mutation occurs exactly once in the phylogeny. As such, each mutation occurs in exactly one of the subtrees corresponding to the $p$ copy-number clusters. We will show frequency $f_{i,j} = 0$ or $f_{i,j} = 1$ if mutation $j$ does not occur in subtree corresponding to copy-number cluster $i$. Let vertex $v_i$ which denotes root of the subtree of copy-number cluster $i$. There are two cases to consider if mutation $j$ does not occur in subtree of cluster $i$. First, mutation $j$ lies on the path from root to vertex $v_i$. As such, clones corresponding to $v_i$ and every vertex downstream of $v_i$ have mutation $j$. Since clone proportions $\sum_\ell u_{i,\ell} = 1$, we must have $f_{i,j} = 1$. Second, mutation $j$ does not lie on path from root to vertex $v_i$. Since mutation $j$ does not occur in the subtree of cluster $i$, none of the clones represented by vertices in the subtree have mutation $j$. Therefore, $f_{i,j} = 0$.

**Condition (2)** Since $F$ is generated by copy-number constrained perfect phylogeny $T$, $u_{i,\ell} > 0$ only if clone $\ell$ corresponds to vertex in subtree corresponding to cluster $i$, i.e. $\sigma(v) = i$. If $f_{i,j} = 1$ and $f_{i,j'} < 1$, then there must be a clone $\ell$ such that $b_{\ell,j} = 1$ and $b_{\ell,j'} = 0$. Therefore $j$ must occur before $j'$ in the phylogeny $T$ and thus by construction we have $j \prec_S j'$.

**Condition (3)** Since $\sum_\ell u_{i,\ell} = 1$, $f_{i,j} = 1$ if and only if all clones with non-zero proportion ($u_{\ell,j} > 0$) in copy-number cluster $i$ have every mutation $j$ where $f_{i,j} = 1$. Since $T$ is a constrained perfect phylogeny, all these clones form a subtree and the root $v_i$ of this subtree must have all these mutation, i.e. these mutation lie on the path from root to $v_i$. Moreover, a mutation $j'$ can't be on this path if $f_{i,j'} < 1$ since all clones will have mutation $j'$ and $\sum_\ell u_{\ell,j'} = 1$.

**Condition (4)** Since $T$ is consistent with forbidden ancestral pairs $D$, there is no pair of copy-number clusters $(i, i') \in R$ such that subtree of $i'$ is nested in subtree of $i$. Suppose condition (4) is not true and there exists a mutation $j$ that is subclonal in cluster $i$ and clonal in cluster $i'$. Then there must exist a vertex $u$ such that $\sigma(u) = i$ and the incoming edge to $u$ indicates gain of mutation $j$. Since $j$ is clonal in cluster $i'$, there must be at least one vertex $u'$ such that $\sigma(u') = i'$ and $u \prec_T u'$. However, this contradicts the premise that $T$ is consistent with $D$.

**Condition (5)** Consider mutation $j$ with children $\delta_S(j)$. Since $T$ is a perfect phylogeny, every clone that has mutation from the set $\delta_S(j)$ must also have mutation $j$. Therefore $f_{i,j} = \sum_\ell u_{\ell,j}\mathbf{1}(b_{\ell,j} = 1) \geq \sum_\ell \sum_{j' \in \delta_S(j)} u_{\ell,j'}\mathbf{1}(b_{\ell,j'} = 1)$. Therefore, $f_{i,j} \geq \sum_{j' \in \delta_S(j)} f_{i,j'}$, which is the sum condition.

($\Leftarrow$) Suppose cell fraction $F$ and forbidden ancestral pair $D$ satisfy the five constraints stated in the theorem for some mutation tree $S$. We will show that there exists a copy-number constrained perfect phylogeny $T$ that is consistent with $D$ and generates cell fraction matrix $F$. We start by proving a few results which will be useful in the construction of $T$.

From condition (3), for each copy-number cluster $i$, there must a vertex $v_i$ in $S$ mutation $j$ is on the path from root of $S$ to $v_i$ if and only if mutation $j$ is clonal in cluster $i$, i.e. $f_{i,j} = 1$. We claim that all mutations that are subclonal in cluster $i$ must form a connected subtree in $S$ rooted at vertex $v_i$. We show this as follows. Due to condition (2), every vertex in $S$ corresponding to mutation $j'$ that is subclonal in cluster $i$ must occur downstream of vertex $v_i$, i.e. $v_i \prec_S j'$. We show that the such vertices also form a connected subtree by contradiction. Suppose there are two vertices $v'$ and $v''$ such that $v' \prec_S v''$, where mutation corresponding to $v''$ is subclonal in cluster $i$ but mutation corresponding to $v'$ is not, i.e. $f_{i,j'} = 0$. However, this contradicts condition (5), that $f_{i,j'} > f_{i,j''}$ for every cluster $i$ if $j' \prec_S j''$.

If two copy-number cluster $i$ and $i'$ have the same set of clonal mutations and just $v_i = v_{i'}$ the connected subtrees formed by the subclonal mutations will be disjoint (not share any vertices) except the roots $v_i$ and $v_{i'}$, respectively. This is because of Condition (1) which states that no two copy-number clusters can share a subclonal mutation.

We obtain such a phylogeny $T$ from $S$ as follows. We start with constructing a tree $T$ with the same topology as $S$ and label each vertex $v$ with a binary vector $b_v \in \{0,1\}^m$, where $b_{v,j} = 1$ if and only if mutation $j$ lies on the path from root to the vertex in $S$. Identify the vertex $v$ for each copy-number cluster $i$ such that $b_{v,j} = 1$ if and only if $f_{i,j} = 1$. We introduce a new vertex $v'$ with edge $(v, v')$ and move the subtree with subclonal mutations of cluster $i$ rooted at $v$ to $v'$. We label the new vertex $v'$ with $\sigma(v') = i$ and $b_{v'} = b_v$. Moreover, each vertex $v$ in the subtree of cluster $i$ is also labeled with $\sigma(v) = i$. The remaining vertices are given a null label ($\sigma(v) = \emptyset$). The resulting tree $T$ by construction satisfies condition (i), (ii) and (iii) of Definition 2.1. We need to show that $T$ satisfies condition (iv), i.e. $T$ is consistent with forbidden ancestral pairs $D$.

We prove consistency of $T$ with $D$ by contradiction. Suppose $T$ is not consistent with $D$, i.e. there is a pair $(i, i')$ of copy-number clusters in $D$ such that there is a pair of vertices $v \prec_T v'$ with $\sigma(v) = i$ and $\sigma(v') = i'$. Observe that these clusters cannot have the same set of clonal mutations, because in that case, by construction, their subtrees with be disjoint such pair $(v, v')$ of vertices would not exist. By construction, it must be the case that mutation $j$ corresponding to vertex $v$ in $S$ precedes mutation $j'$ corresponding to vertex $v'$ in $S$, i.e. $j \prec_S j'$. Since $\sigma(v) = i$, mutation $j$ must be subclonal in cluster $i$ and since $j \prec_S j'$ it must be clonal in cluster $i'$, which contradicts Condition (4) of Theorem 4.2. This concludes the proof.

□

We restate Corollary 4.1 here and provide a proof.

**Corollary A.1.** *If a copy-number constrained perfect phylogeny $T$ generates a cell fraction matrix $F$, then its binarization $F'$ is a perfect phylogeny matrix.*

*Proof.* Let $F$ be a $p \times m$ cell fraction matrix generated by a copy-number constrained perfect phylogeny $T$. Then by Theorem 4.2, there is a mutation tree $S$ on the $m$ mutations where for every row $i$, there exists a vertex $v$ in $S$ such that mutation $j$ is on the path from root to $v$ in $S$ if and only if $f_{i,j} = 1$. We will show that binarization $F'$ of $F$ is a perfect phylogeny matrix, i.e. it admits a perfect phylogeny.

There are several equivalent characterizations of perfect phylogeny matrices [52, 53]. We will use the following characterization: binary matrix $A$ is a perfect phylogeny matrix if and only if for any two columns, the sets of rows with 1 entries are either disjoint or related by containment. It is easy to show that $F'$ has this property. The columns are $F'$ correspond to mutations (vertices of the mutation tree). Consider two mutations $j$ and $j'$. In the mutation tree $S$, the vertices $v_j$ and $v_{j'}$ for these mutations would have one of three relation: (i) $v_j$ and $v_{j'}$ are incomprable, i.e. on distinct branches of $S$, (ii) $v_j \prec_S v_{j'}$ and (iii) $v_{j'} \prec_S v_j$. In the first case, the set of vertices for which mutation $j$ lies on the path from root to that vertex and the set of vertices for which mutation $j'$ lies on the path from root to that vertex, will be disjoint. As such, for these two columns in $F'$, the sets of rows with 1 entries will be disjoint. In the second and third case, the set of vertices for which mutation $j$ (or $j'$) lies on the path root to that vertex will be a superset of the set of vertices for which mutation $j'$ (or $j$) lies on the path from root to that vertex. As such, for these two columns in $F'$, the sets of rows with 1 entries will be related by containment. This concludes the proof. □

##### A.2 Proof to Theorem 4.3

We restate the theorem here for completeness and provide a proof in the following.

**Problem A.1.** *[Decision Version of Constrained Perfect Phylogeny Mixture with Errors (dCPPM-E)] Given matrices with cell-fraction confidence intervals $F^-, F^+ \in \mathbb{R}^{n \times m}$ for $n$ cells and $m$ mutations, forbidden ancestral pairs $D$, and an integer $k \geq 0$, does there exist a subset of mutations $S \subseteq \{1, \ldots m\}$ with $|S| \geq m - k$ such that for the* *submatrices, $F'^-$, $F'^+$, of $F^-$, $F^+$ restricted to the columns in S, there is a cell-fraction matrix $F \in \mathbb{R}^{n\times|S|}$, tumor phylogeny T, clone proportion matrix U, mutation matrix B and copy number clustering $\sigma$ that satisfies the following conditions: (i) $f'^-_{i,j} \le f_{i,j} \le f'^+_{i,j}$ for each cluster i and mutation $j \in S$, (ii) $F = UB$, (iii) T is a constrained perfect phylogeny for B and $\sigma$ that is consistent with D, and (iv) $u_{i,\ell} = 0$ if $\sigma(\ell) \neq i$.*

**Problem A.2.** *[Decision Version of Minimum Character Removal Perfect Phylogeny Problem (dMCR)] Given a binary matrix $M \in \{0,1\}^{n\times m}$ whose rows represent taxa and columns represent binary characters, and an integer $k \ge 0$, does there exist a $S \subseteq \{1,\ldots,m\}$ with $|S| \ge m - k$ such that the submatrix of M restricted to the columns in S satisfies the three-gamete condition, and therefore admits a perfect phylogeny?*

This problem is known to be NP-hard [54].

**Theorem A.2.** *Decision version of the CPPM-E problem is NP-hard.*

*Proof.* We show this by constructing a polynomial-time reduction from the decision version of dMCR (Problem A.2) to the decision version of CPPM-E (dCPPM-E) (Problem A.1).

Let an instance of dMCR be a binary matrix $M \in \{0,1\}^{n\times m}$ and an integer $k \ge 0$. We produce an instance of dCPPM-E as follows.

1. We use the same numbers for $n$ (clusters / rows) and $m$ (mutations / columns).
2. We set the cell-fraction interval matrices elementwise to the exact values of $M$: $F^- = F^+ = M$, i.e, i.e. for every $i, j$ we set $f^-_{i,j} = f^+_{i,j} = x_{i,j}$.
3. We let the allowed number of removed mutations be the same $k$.
4. We set the number of clones $r = n$ and fix a cluster assignment $\sigma$ that assigns clone $\ell$ to cluster $\ell$. So, $\sigma(\ell) = \ell$ for $\ell = 1,\ldots,n$. We set no forbidden pairs, i.e, $D = \emptyset$.

This construction takes $O(nm)$ i.e., polynomial time.

We now prove correctness (equivalence of true-instances).

($\Rightarrow$) Suppose the dMCR instance has a solution: there exists $S \subseteq \{1,\ldots,m\}$ with $|S| \ge m-k$ such that the submatrix $M'$ of $M$ with columns in $S$ satisfies the three-gamete condition (and therefore admits a perfect phylogeny).

We consider the corresponding dCPPM-E instance restricted to the same column set $S$. We choose $F = M', U = I_n, B = M'$. Then $F^-_{:,S} \le F \le F^+_{:,S}$ holds, and in fact they are equal [condition (i)]. Also $F = UB$ [condition (ii)], and $B = M'$ admits a perfect phylogeny $T$. Since, every node in $T$ has its own cluster label, the clusters from separate subtrees with one node each. Since, $D =, T$ trivially fulfills the forbidden pair constraints as well. Hence, according to Defintion 2.1 $T$ is a copy-number constrained phylogeny tree [condition (iii)]. With $\sigma(\ell) = \ell$ and $U = I$, we have $u_{i,\ell} = 0$ whenever $\sigma(\ell) \neq i$ [condition (iv)].

Thus the dCPPM-E instance has a feasible solution for the same set $S$.

($\Leftarrow$) Conversely, we suppose the constructed dCPPM-E instance is a true-instance: there exists $S$ with $|S| \ge m - k$ and matrices $F$, $U$, $B$, a tree $T$, and mapping $\sigma$ satisfying the dCPPM-E conditions on columns $S$.

Since $F^- = F^+ = M$, the interval constraint forces $F$ to equal the submatrix $M'$ of $M$ on columns $S$; that is, $F = M'$. Condition (ii) then yields $M' = UB$. According to the construction $U = I_n$. Hence $M' = B$. Condition (iii) states that $B$ admits a (constrained) perfect phylogeny $T$ for the given $\sigma$. Because $B$ admits a perfect-phylogeny labeling for exactly the characters (columns) in $M'$, the matrix $M'$ must itself admit a perfect phylogeny. Therefore $M'$ is a valid solution to the original MCR instance.

Since the reduction is polynomial-time and the two instances are equivalent, dCPPM-E is NP-hard because MCR is NP-hard [54]. Since the decision version of CPPM-E is NP-hard, the optimization version must also be NP-hard. ☐

### B Algorithmic details of SCOPE

Here, we provide algorithmic details of SCOPE regarding: (1) estimation of cell fractions and mutation groups from scWGS data, (2) mixed integer linear program to enumerate all largest constrained perfect phylogeny consistent with estimated cell fractions and forbidden ancestral copy-number cluster pairs, (3) ranking the multiple phylogenies consistent with the data, and (4) placing unassigned mutations on the inferred phylogeny.

##### B.1 Estimating cell fractions and mutation clusters from scWGS data

We sum the read counts and variant counts for each mutation $j$ to get total read count $r_{i,j}$ and variant read counts $q_{i,j}$ for each CN cluster $i$ and mutation $j$. The Variant Allele Frequency (VAF), $v_{i,j}$ of mutation $j$ in CN cluster $i$ is then $v_{i,j} = q_{i,j}/r_{i,j}$. Let $c_{i,j}$ be the copy number at the locus of mutation $j$ for cells in copy number cluster $i$. Within a copy number cluster $i$, for each mutation $j$, the mutated cells can have exactly $x_{i,j}$ mutated copies. This is because under the perfect phylogeny assumption, unmutated copies of the locus cannot gain mutation $j$ and mutated copies cannot be lost. Since all cells within a copy number cluster $i$ share the same copy number $c_{i,j}$ at the mutation locus, the number of mutated copies $x_{i,j}$ must be identical across all mutated cells in that cluster. Hence, this removes all possibilities of varying $x_{i,j}$ for mutated cells within a copy-number cluster.

Therefore, the VAF of mutation $j$ within copy-number cluster $i$ is $v_{i,j} = \frac{x_{i,j}}{c_{i,j}} f_{i,j}$. Hence, $f_{i,j} = \frac{v_{i,j} c_{i,j}}{x_{i,j}}$.

**Case 1:** if mutation $j$ is *gained* inside copy-number cluster $i$, then $x_{i,j} = 1$, and $f_{i,j} = v_{i,j} c_{i,j}$.

**Case 2:** if mutation $j$ is *clonal* to copy-number cluster $i$, then $x_{i,j} \geq 1$, $f_{i,j} = 1$ and $f_{i,j} \leq v_{i,j} c_{i,j}$ which implies $v_{i,j} c_{i,j} \geq 1$.

**Case 3:** if mutation $j$ is *non-clonal* to copy-number cluster $i$, then $f_{i,j}$ then $f_{i,j} = 0$, and, $v_{i,j}$ = 0.

Hence, the cell fraction $f_{i,j}$ for each mutation $j$ in cluster $i$ can be estimated from $v_{i,j} c_{i,j}$. When, $v_{i,j} c_{i,j} < 1$, estimated value of $f_{i,j} = v_{i,j} c_{i,j}$. When, $v_{i,j} c_{i,j} \geq 1$, the estimated value is $f_{i,j} = 1$. Thus, the estimated cell fraction can be expressed compactly as $f_{i,j} = \min(v_{i,j} c_{i,j}, 1)$.

Mutations that have similar copy-number cluster specific cell fractions are clustered together into mutation groups. The cell fractions of individual cells may have errors due to low cumulative read counts $r_{ij}$. Grouping mutations together gives a better estimate of the cell fractions. We use $k$-means clustering using cell fractions as features to group together mutations. We run $k$-means clustering with $k$ values in $[7, 20]$ with 10 random restarts for each and pick the clusterings with the low silhouette scores. We use the upper and lower quantiles of the cell fractions of each mutation group for each copy-number clusters as the confidence interval bounds $F^+$ and $F^-$.

##### B.2 MILP formulation

In the following we show the constraints that enforce conditions (1), (2), (3), (4) and (5) of Theorem 4.2. For ease of exposition, we start with subclonality condition (condition (1)) and the sum condition (condition (5)), followed by clonal consistency of copy-number clones (condition (3)), subclonal mutation condition (condition (2)) and forbidden ancestral pair condition (condition (4)). We end this section with a description of our iterative approach to enumerate all solutions to the CPPM-E problem.

The frequency matrix is encoded by continuous variables $f_{i,j}$ for copy-number cluster $i$ and mutation $j$. We also introduce a variable $x_j$ for each mutation $j \in \{1, \ldots, m\}$ which indicates if a mutation is selected in the solution. If a mutation $j$ is selected, we require that the frequency is bounded from below by $f^-_{i,j}$ and from above by $f^+_{i,j}$. We enforce this with the following constraints for all clusters $i$ and mutations $j$,

$$f_{i,j} \geq x_j f^-_{i,j}, \quad f_{i,j} \leq x_j f^+_{i,j}.$$

The above constraints also remove mutations that are not selected ($x_j = 0$) from the solution by setting corresponding columns in the frequency matrix to 0.

We introduce binary variables $g_{i,j}$ to indicate if mutation $j$ is subclonal in copy-number cluster $i$. Since every selected mutation (i.e. when $x_j$) can be subclonal in at most one copy-number cluster (Condition 1 of Theorem 4.2), for each mutation $j$ we enforce,

$$\sum_{i=1}^{p} g_{i,j} \leq p - (p-1)x_j,$$

and we enforce $0 < f_{i,j} < 1$ for a mutation is subclonal using the following constrains for each mutation $j$ and cluster $i$,

$$\begin{cases} g_{i,j} \geq f_{i,j}, & \text{if } f^-_{i,j} = 0 \text{ and } f^+_{i,j} < 1, \\ g_{i,j} = 1, & \text{if } 0 < f^-_{i,j} < f^+_{i,j} < 1, \\ g_{i,j} \geq 1 - f_{i,j}, & \text{if } f^-_{i,j} > 0 \text{ and } f^+_{i,j} = 1. \end{cases}$$

##### B.2.1 Sum condition

El-Kebir et al. [39] showed that sum condition is satisfied for some mutation tree $S$ if and only if there exists a perfect phylogeny matrix $B$ such that $F = UB$.

We introduce binary variables $b_{\ell,j}$ for each clone $\ell \in \{1, \ldots, r\}$ and mutation $j$ to indicate the presence/absence of mutations in the clones. Clone proportions are encoded using continuous variable $u_{i,\ell}$ for each copy-number cluster $i$ and clone $\ell$ along with constraints $\sum_{\ell=1}^{r} u_{i,\ell} = 1$. We encode the sum condition $F = UB$ by introducing continuous $a_{i,j,\ell}$ to linearize the product $u_{i,\ell} b_{\ell,j}$ using the following constraints for each copy-number cluster $i$, clone $\ell$ and mutation $j$,

$$a_{i,j,\ell} \leq u_{i,\ell}, \quad a_{i,j,\ell} \leq b_{\ell,j}, \quad a_{i,j,\ell} \geq u_{i,\ell} + b_{\ell,j} - 1.$$

Enforcing $F = UB$ is achieved by adding the constraint $f_{i,j} = \sum_{\ell=1}^{r} a_{i,j,\ell}$ for each copy-number cluster $i$ and mutation $j$. Perfect phylogeny constraints on $B$ along with clonal consistency are shown in the next section.

##### B.2.2 Perfect phylogeny constraints

We enforce $B$ to be a perfect phylogeny matrix and impose clonal consistency of copy-number clusters by enforcing that the stacked matrix $[B; F']$ is a perfect phylogeny matrix. We encode the stacked matrix $[B; F']$ by introducing variables $b_{\ell,j}$ for $\ell \in \{r+1, \ldots, r+p\}$, representing the rows of $F'$ and impose the following constraints for all copy-number clusters $i$ and mutations $j$,

$$b_{n+i,j} \leq f_{i,j}, \;\; b_{n+i,j} \leq 1 + g_{i,j}, \;\; b_{n+i,j} \geq f_{i,j} - g_{i,j}.$$

We enforce that the extended matrix $B \in [0,1]^{(r+p)\times m}$ is a perfect phylogeny using the set inclusion and disjointness (SID) formulation described in Chimani *et al.* [53]. This formulation uses a characterization of perfect phylogeny matrices that states that for any two columns, we require the 1-sets, i.e. set of rows $i$ such that $b_{\ell,j} = 1$, of any two columns should either be disjoint or related by containment [52]. We introduce continuous variables $y_{j,j'}$ and $z_{j,j'}$ for each pair of mutations $j$ and $j'$. We force $y_{j,j'} = 0$ if the 1-set of mutation $j$ is not contained in 1-set of mutation $j'$ using the following constraint for each row $\ell$,

$$y_{j,j'} \leq 1 - b_{\ell,j} + b_{\ell,j'}.$$

Along the same vein, we enforce $z_{j,j'} = 0$ if the 1-set of mutation $j$ is not disjoint with 1-set of mutation $j'$ using the following constraint for each row $\ell$,

$$z_{j,j'} \leq 2 - b_{\ell,j} - b_{\ell,j'}.$$

For $B$ to be a perfect phylogeny matrix, we require that for any two mutations $j$ and $j'$, the 1-sets should be either related by containment or be disjoint. In other words, at least one of $y_{j,j'}$, $y_{j',j}$ and $z_{j,j'}$ must be 1 for each pair of mutation $j$ and $j'$. We achieve this by enforcing the following constraint for each pair $j, j'$ of mutations,

$$y_{j,j'} + y_{j',j} + z_{j,j'} \geq 1.$$

In summary, to enforce that $B$ is a perfect phylogeny matrix we introduce $O(m^2)$ continuous variables and $O(nm^2 + pm^2)$ constraints.

##### B.2.3 Consistency between clonal and subclonal mutations in copy-number clusters

According to Condition (2), we require that every subclonal mutation $j$ of a copy-number cluster $i$ but be descendant of a clonal mutation $j'$ of cluster $i$. We enforce this by requiring that $y_{j,j'}$, which is forced to 0 if $j$ is not descendant of $j'$ in the perfect phylogeny constraints described above, must be 1 if $j'$ is clonal ($b_{i,j'} = 1$) and $j'$ is subclonal ($g_{i,j} = 1$) for a copy-number cluster $i$, using the following constraint,

$$y_{j,j'} \geq b_{i,j'} + g_{i,j} - 1.$$

We impose the above constraint for every pair of mutations $j, j'$ and every copy-number cluster $i \in [p]$.

##### B.2.4 Enforcing uniqueness of solutions

Since we want to generate all unique solutions, we must ensure that we do not repeated solution where the rows of mutation matrix $B$ are permuted. As such, as enforcing symmetry breaking constraints to enforcing an ordering on the rows of $B$ as follows,

$$\sum_{j=1}^{m} 2^j b_{\ell,j} \leq \sum_{j=1}^{m} 2^j b_{\ell+1,j} - 1, \quad \text{for all } \ell \in \{1, \ldots, r-1\}.$$

##### B.2.5 Incorporating forbidden ancestral copy-number clusters

To enforce that copy-number cluster $i'$ is not descendant of copy-number cluster $i$, it suffices to ensure that any mutation $j$ gained in cluster $i$ (i.e. $g_{i,j} = 1$) is not clonal in cluster $i'$ (i.e. $b_{i',j} = 1$). As such for every forbidden ancestral pair $(i, i')$ in the input, we enforce the following constraints,

$$g_{i,j} + b_{i',j} \leq 1, \quad \text{for all } j \in \{1, \ldots, m\}.$$

##### B.2.6 Objective function

We set $\sum_{j=1}^{m} x_j$ as the objective function to maximize the number of selected mutations. The complete MILP has $O((n+p)m + m^2)$ binary variables, $O(mnp + pm + pn)$ continuous variables, and $O(mnp + (n+p)m^2 + |D|m)$ constraints.

##### B.2.7 Enumerating all solutions

For a given confidence interval $F^-$ and $F^+$, any feasible solution of CPPM-E must satisfy,

$$F = UB, \quad F^-_{i,j} \leq F_{i,j} \leq F^+_{i,j}$$

Note that $U$ and $F$ are continuous real variables and so even when the mutation matrix $B$ is fixed, there may be infinitely many choices of $U$, and therefore $F$, that satisfy the interval constraints. However, such variations in $U$ and $F$ alone do not change the underlying phylogeny tree, defined by $B$, and therefore are not of interest to us. Thus, for CPPM-E we enumerate solutions with *distinct* $B$ matrices.

To guarantee that each newly found solution differs from every previously discovered solution, we add a distinction constraint for each previously found mutation matrix $B^{(k)}$. To forbid $B^{(k)}$, we require that at least one entry of $B$ must differ from the corresponding entry of $B^{(k)}$:

$$\sum_{i,j} |b_{i,j} - B^{(k)}_{i,j}| \ \geq \ 1.$$

This constraint forces the ILP to exclude the exact binary pattern $B^{(k)}$ from future solutions.

The absolute value is linearizing the constraint as:

$$\sum_{i,j} \begin{cases} 1 - b_{i,j}, & B^{(k)}_{i,j} = 1, \\ b_{i,j}, & B^{(k)}_{i,j} = 0 \end{cases} \ \geq \ 1$$

This ensures that the new solution must differ from every previously found $B^{(k)}$ in at least one entry. We add such constraint for all previously found solutions when finding a new solution. This adds $\Theta((n+p)m)$ constraints for each new solution stored.

If we require *distinct mutation trees only*, we apply the same distinct constraint but restrict the summation to the rows of $B$ that encode clone-mutation assignments. In this case, two solutions are considered distinct only if their mutation trees differ, even if rows for the copy-number clusters in $B$ differ.

##### B.3 Ranking solutions based on deviation from median cell fractions

After enumerating feasible mutation matrices $B$ that satisfy the CPPM-E constraints, each $B$ defines a family of admissible decompositions $(U, F)$ such that $F \in [F^-, F^+]$ and $F = UB$, where $F^-$ and $F^+$ denote the lower and upper confidence bounds for the cell fractions. For a fixed phylogeny tree (equivalently, a fixed mutation matrix $B$), the remaining degrees of freedom lie in the continuous variables of the clone proportion matrix $U$ and the realized cell fraction matrix $F$. Different $(U, F)$ pairs corresponding to the same $B$ represent alternative decompositions, but some decompositions adhere to the empirical estimation of cell fractions more closely than others. To prioritize among these decompositions, we re-solve the ILP for each enumerated $B$ while fixing the mutation matrix $B$ and optimizing only the continuous variables $U$ and $F$. Specifically, we introduce an objective that penalizes deviation of the realized cell fractions from their empirical medians. Let $\tilde{F}$ denote the matrix of observed median cell fractions for mutation groups. We then solve

$$\min_{U,F} \sum_{s,\ell} \left| F_{s\ell} - \tilde{F}_{s\ell} \right| \tag{1}$$

subject to

$$F^- \leq F \leq F^+, \quad F = UB, \quad U \geq 0, \quad \sum_{\ell} u_{i,\ell} = 1, .$$

This refinement step selects, for each candidate mutation tree, the decomposition that best matches the central tendency of the observed data. Hence, for each mutation matrix $B$ or phylogeny tree we get the $U$ and $F$ matrices with the closest fit to the data and also the corresponding least deviation from the data.

For the same mutation tree, there can be multiple cluster placements. For a mutation tree, we can choose the cluster placement that has the best fit to data or the lowest deviation. We do this by optimizing the same optimization objective as 1, subject to copy-number constrained perfect phylogeny constraints (Theorem 4.2, formulated as MILP in Sec. B.2). In this way, each mutation tree is associated with its optimal cluster placement and its minimal achievable deviation.

We can rank the phylogeny tree or mutation tree solutions by the order of lowest deviation from data. This prioritizes trees with the closest fit to the data.

##### B.4 Placing unassigned mutations on the phylogeny

Due to the sparsity in ultra-low coverage scWGS data, there is a lot of uncertainty in the mutation groups. This may lead to a lot of mutation being unassigned in the phylogeny, since we select mutations in groups in the MILP. To address this, we employ a heuristic to place unassigned mutations on the phylogeny produced by the MILP.

Specifically, we calculate the Euclidean distance between each unassigned mutation's cell-fraction vector and the median profile of each selected mutation group, and transform these distances into normalized responsibility scores using a temperature-scaled softmax, such that higher values indicate better matches. Each mutation is assigned to the mutation group with the highest responsibility provided that this value exceeds a predefined confidence threshold ($0.2$ in our analyses), ensuring that assignments are made only when there is sufficient support. Mutations that do not meet this criterion remain unassigned. This procedure allows us to incorporate previously unassigned mutations into the phylogeny in a principled and reliable way, minimizing the risk of introducing errors.

#### C Simulation details

We generated single-cell whole genome sequencing data by simulating the evolutionary processes of tumor heterogeneity: the hierarchical acquisition of somatic mutations and copy number alterations, followed by noisy observation through sequencing technology. We used these data to compare SCOPE with other tumor phylogeny inference methods. We generate a phylogeny $T$ over $m$ mutation groups and $p$ copy number clusters, with $m \in \{5, 10, 15\}$ and

$p \in \{5, 10, 15\}$. The mutation groups and copy number clusters mark clonal expansion of tumor events: somatic mutations and copy number aberrations. The tumor events occur in uniform random temporal order. The tree topology is formed using tumor events as nodes through a random attachment process. A mutation group is subclonal to its nearest ancestor copy number cluster node.

Each mutation group contains a variable number of individual mutations, with group sizes drawn from a Poisson distribution with mean $S \in \{100, 500, 1000\}$. The genome is divided into 1000 genomic bins, and each mutation is randomly assigned to one of these bins. Each copy number cluster is characterized by the copy numbers of these 1000 bins. The copy numbers are sampled uniformly from $\{1, \ldots, 8\}$ for each cluster-bin pair. Each mutation belongs to the copy number cluster to which its mutation group is subclonal to. The copy number of a mutation is determined by the genomic bin in which it is assigned to and the copy number cluster to which it belongs. We impose no mutation losses, ensuring all mutations follow perfect phylogeny on the mutation tree.

We simulate $n \in \{1000, 5000, 10000\}$ cells by randomly assigning cells to tree nodes. A cell has all the mutations of its mutation group and their ancestor groups. The copy number cluster of a cell is that of its mutation group. The total read counts $N_{i,j}$ of each cell $i$ for each mutation $j$ are drawn from a Poisson distribution with mean $cov \in \{0.02, 0.05, 0.1\}$, modeling the ultra-low coverage of whole genome single-cell data. The latent true variant allele fraction VAF, $f_{i,j}$ of a cell $i$ for a mutation $j$ present in bin $k$ is $1/$copy number of bin $k$. If mutation $j$ is absent in cell $i$, then $f_{i,j} = 0$. We incorporate systematic errors with false positive rate $\alpha = 0.001$ and false negative rate $\beta = 0.001$, for which we get an error-corrupted VAF of $f'_{i,j} = \alpha + (1 - \alpha - \beta) f_{i,j}$. Variant read counts follow a beta-binomial distribution: $V_{i,j} \sim \mathrm{BetaBinom}(N_{i,j}, f'_{i,j}\gamma, (1 - f'_{i,j})\gamma)$, where $\gamma = 15$ corresponds to overdispersion from allelic dropout and amplification bias.

We produce data for different values of parameters: $n$, $m$, $p$, $S$, $cov$, yielding 243 distinct parameter combinations with 5 independent replicates each for a total of 1,215 datasets.

#### C.1 Performance Evaluation with Competing Methods

##### C.1.1 Pairwise ancestral relationship accuracy

Here, we describe the computation of pairwise ancestral relation accuracy $E(T, T')$ for two tumor phylogenies $T$ and $T'$. Under the assumption that a mutation can be gained only on in the phylogeny, any pair $(j, j')$ of mutations can be related in exactly one of the following four ways.

1. Mutation group of mutation $j$ that is an ancestor of the mutation group of mutation occurs along the path from the root to source node of edge on which mutation $j'$ occurs.
2. Mutation group of mutation $j$ that is an descendant of the mutation group of mutation occurs along the path from the root to source node of edge on which mutation $j'$ occurs.
3. Mutation $j$ and $j'$ are part of the same mutation group.
4. Mutation $j$ and $j'$ are disjoint. They are neither ancestral to each other and nor part of the same mutation group.

We compute the accuracy of inferring the correct relationship between all possible pairs of mutations from the inferred tumor phylogeny.

##### C.1.2 Mutation Placement accuracy

Here, we describe the computation of the mutation placement accuracy $\epsilon(T, T')$ for two tumor phylogenies $T$ and $T'$. We are assuming that a mutation can be gained only once in a phylogeny, and hence, it can be gained within at most one copy number cluster. We call the copy number cluster in which a mutation has been gained as the cluster placement of the mutation. For each mutation, we compute the accuracy of inferring the correct cluster placement.

SCOPE gives a phylogeny tree with nodes showing both copy number change events and SNV mutation events. As a result, the cluster placement of a mutation is simply the nearest ancestor copy number change event in the phylogeny tree.

**Computing mutation placement accuracy for Phertilizer** Phertilizer produces clone trees and assigns cell and SNV to the clones on the tree, but does not place copy-number (CN) events on the tree. We first infer where CN cluster should lie on Phertilizer 's tree by using the clone assignments of individual cells. The distribution of ground-truth CN clusters of the cells of a predicted clone defines that clone's CN-cluster composition. For each CN cluster, we identify the subtree rooted at the the lowest common ancestor (LCA) of all clones containing cells from that cluster. Any mutation gained within this subtree is *placed* in that copy-number cluster. According to this procedure, a mutation may be placed in multiple copy-number clusters. However, in the ground-truth phylogeny which follows the copy-number constrained perfect phylogeny model, each mutation is placed in a unique copy-number cluster. A mutation's accuracy in the Phertilizer phylogeny is defined as the fraction of its predicted CN clusters that match its true CN cluster (with zero assigned if no clusters are predicted); the overall cluster-placement accuracy of the phylogeny is the average of these per-mutation accuracies.

##### C.2 Generating Phylogeny Tree from SBMClone Clusters

SBMClone gives as output a clone $\times$ mutation cluster matrix $M$. The value $M_{i,j}$ in the matrix represents the proportion of cells in the clone $i$ having mutations in mutation group $j$. The matrix $M$ is binarized into $M'$ as follows: $M'_{i,j} = 1$ if $M_{i,j} \geq \tau$, for a threshold $\tau$. The value $M'_{i,j}$ in the matrix tells if clone $i$ has mutation group $j$ . With this binarized matrix $M'$ we construct the phylogeny tree under the perfect phylogeny model [52]. We used a threshold of $\tau = 0.01$.

#### D Mutation clustering and forbidden ancestral pairs in real data

We processed the real datasets by first generating cluster-specific cell fractions for each mutation, using the procedure described in B.1, based on mutation-specific total read counts, variant counts, and CN cluster assignments. We then grouped mutations using $k$-means clustering.

For Laks *et al.* OV2295 dataset [8] we did $k$-means clustering with $k \in [7, 20]$ and 10 random restarts for each value of $k$. Clustering quality was evaluated using silhouette scores. Although no clustering showed a clear elbow, $k = 12, 17, 20$, and 24 showed potential inflection points. We selected these $k$ values and run SCOPE on all of these. For each cluster, we got the cluster-specific cell-fraction intervals for each mutation group using the 25th and 75th percentiles of the cell fractions of the mutations in the group. SCOPE was run using the mutation clusterings using these cell fractions of mutation groups as input.

SCOPE was run using cell fraction of mutation groups obtained from clustering with $k = \{12, 17, 20, 24\}$ mutation groups. Finally, we chose $k = 17$ mutation groups because it placed the largest number of mutations into the inferred phylogeny.

For all the meta-cohort samples [19], did $k$-means with $k \in [15, 20]$. We selected the clustering with the lowest silhouette score and ran SCOPE with the cell fractions of those mutational groups.

To further constrain the phylogeny, loss-of-heterozygosity (LOH) events were used to identify forbidden ancestral copy-number cluster pairs. For each copy-number cluster pair $(i, i')$, we count the number of genomic loci where cluster $i$ retained a non-zero copy number (i.e., no LOH) while cluster $i'$ exhibited LOH (i.e. zero copy number for one of the allele) (Fig. S1). In order to account for errors in the inferred copy-number profiles, we consider $(i, i')$ as a forbidden pair if this count exceeds a fixed threshold. For Laks *et al.* OV2295 dataset [8], we used a threshold of 50. For the meta-cohort samples [19], we used a higher threshold of 100.

Let $c_{i,l}$ denote the copy numbers of at genomic locus $l \in \mathcal{L}$ in an allele, for copy-number cluster $i$. A loss-of-heterozygosity (LOH) event at locus $l$ for an allele is defined as

$$\mathrm{LOH}_{i,l} = \begin{cases} 1, & \text{if } c_{i,l} = 0, \\ 0, & \text{otherwise.} \end{cases}$$

For a pair of clusters $(i, i')$, we define the LOH conflict count as

$$\mathrm{conflict}(i, i') = \sum_{l \in \mathcal{L}} \mathrm{LOH}_{i,l} \big(1 - \mathrm{LOH}_{i',l}\big),$$

which counts the number of loci where cluster $i$ has a LOH while cluster $i'$ retains the loci (no LOH).

A cluster pair $(i, i')$ is considered a forbidden ancestral pair if

$$\mathrm{conflict}(i, i') > \tau,$$

where $\tau$ is a fixed threshold to account for possible errors in the inferred copy-number profiles.

Since, the major allele in the OV2295 dataset [8] did not have any conflicts, we only considered the conflict counts from the minor allele (Fig. S1). We used the threshold of $\tau = 50$ to get the forbidden pairs. For the meta-cohort samples [19] we considered both alleles and used $\tau = 100$.

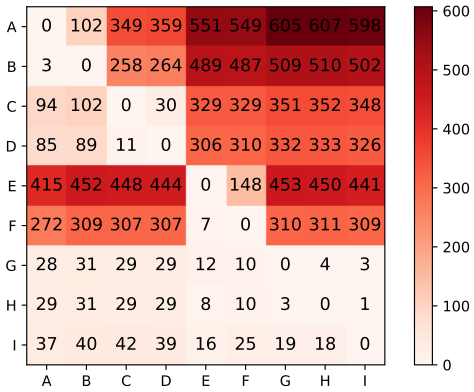

***Fig. S1*** **LOH Conflict Counts for Ovarian Cancer Dataset:** Cell $(i, i')$ has the value of $\text{conflict}(i, i') = \sum_{l \in \mathcal{L}} \text{LOH}_{i,l}\big(1 - \text{LOH}_{i',l}\big)$.

#### E Statistical test for evaluating clonal relationships

We evaluated the clonal relationships between mutation and copy number clusters that we get from the phylogeny trees from different methods. If a mutation group is clonal to a copy number cluster, the cells in that cluster should have the mutations in the group. To test this, we assess whether the observed variant read counts in each copy-number state are compatible with the mutation group being clonal in that cluster.

Let us consider cells in a copy number cluster with copy number state $c$. If a mutation is present in those cells, each sequencing read should be a variant read with probability $p = 1/c$. If we incorporate a sequencing false negative error rate $\varepsilon$, we define an effective probability of getting a variant read as $p_{\text{eff}} = (1 - \varepsilon)/c$. Taking into account for over-dispersion that arise from allelic imbalance and technical variability, the number of variant reads $V$ out of $t$ total reads is assumed to follow a beta-binomial distribution, $V \sim \text{BetaBinomial}(t, \alpha, \beta)$, where the parameters are expressed using a mean-precision parameterization: $\alpha = p_{\text{eff}}\,\theta$, $\beta = (1 - p_{\text{eff}})\,\theta$. Here, $\theta > 0$ controls the degree of over-dispersion.

To test whether the observed variant count $v$ is *too low* to be consistent with the mutation of a mutation group being present in the cluster, we compute the one-sided p-value

$$P(V \leq v \mid t, c) = F_{\text{BB}}(v; t, \alpha, \beta),$$

where $F_{\text{BB}}$ is the beta-binomial cumulative distribution function. Small p-values indicate that the observed variant allele count is lower than expected under the null, suggesting that the mutation may not be present in cells with copy number $c$.

CN cluster may include multiple distinct copy-number states. For each mutation group predicted to be clonal to a CN cluster, we compute a p-value for every CN state represented in that cluster. These p-values are then aggregated using a weighted Stouffer method [49]. The $p$-values are weighed by total read counts in each copy number state.

The resulting combined $p$-value reflects the likelihood of the mutational cluster being clonal in the copy number cluster. Small combined $p$-values indicate that the variant allele read counts are inconsistent with the mutation being clonal in the cluster, thereby providing statistical evidence against specific clonal relationships in the inferred phylogeny. For all tests, we used the parameters $\varepsilon = 0.01$ and $\theta = 300$.

#### F Annotating Mutated Genes

We annotated each mutation group using cancer gene-level information provided by Funnell *et al.* [19] which was derived from the OncoKB [47] database in their analyses. For every clone (or mutational group), we first identified the cancer-relevant genes harboring the largest number of mutations within that group. Among these candidate genes,

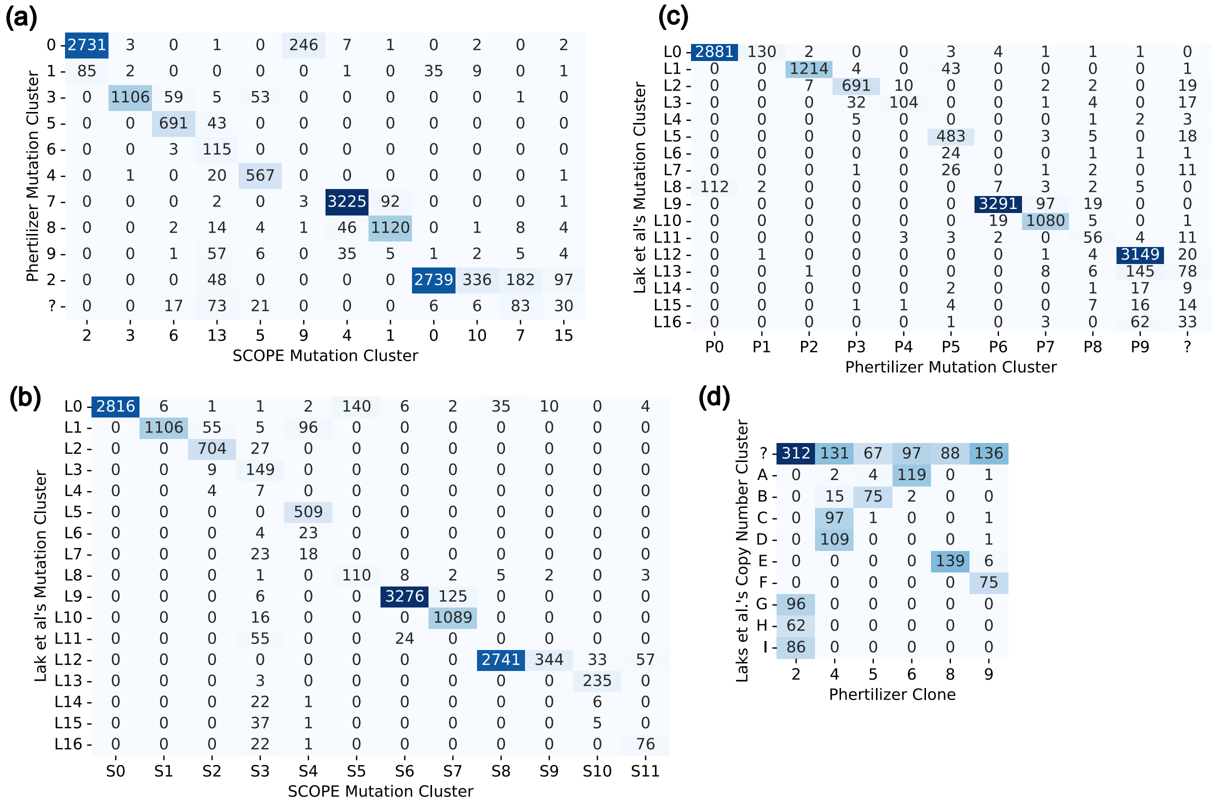

***Fig. S2*** Confusion map showing number of mutations in mutation groups in (a) SCOPE and Phertilizer (b) SCOPE and Laks *et al.* (c) Phertilizer and Laks *et al.* (d) Consistency between copy-number clusters derived by Laks et al. and cell clusters inferred by Phertilizer.

we then selected those with the highest observed mutation frequencies in ovarian cancer cohorts on cBioPortal [48] to assign as the group's representative annotations.

#### G Method parameters

In this section, we provide the commands used to run competing methods for benchmarking on the simulated instances.

##### Phertilizer

We ran Phertilizer with maximum number of iterations set to 10 and with 4 restarts.

```
phertilizer -f {snv_counts_file} --bin_count_data {bin_count_file}
    --tree {tree_output_file} --tree_text {tree_text_output_file}
    -n {cell_clusters_output_file} -m {snv_clusters_output_file}
    -j 10 -s 4
```

##### SBMClone

We ran SBMClone using the following command.

```
python sbmclone.py {input_matrix.csv}
```

#### H Supplementary results

| Cancer Type | Patient | # Mutations | # Cells | # Solutions | # Mutations Selected | # CN Clusters | Coverage | #Forbidden Ancestral Pairs |
|---|---|---|---|---|---|---|---|---|
| HGSOC FBI | SA1091 | 8928 | 506 | 6 | 5943 | 5 | 0.086 | 6/25 |
| | SA1162 | 28171 | 254 | 1 | 13132 | 8 | 0.037 | 38/64 |
| | SA1180 | 6138 | 774 | 1 | 2757 | 13 | 0.074 | 87/169 |
| HGSOC HRD-Dup | SA1050 | 27266 | 990 | 6 | 15525 | 8 | 0.068 | 38/64 |
| | SA1052 | 38881 | 556 | 4 | 25115 | 8 | 0.064 | 49/64 |
| | SA1181 | 10036 | 296 | 1 | 4507 | 6 | 0.119 | 6/36 |
| | SA1184 | 5816 | 621 | 4 | 4067 | 7 | 0.078 | 21/49 |
| HGSOC TD | SA1093 | 16051 | 346 | 1 | 9674 | 4 | 0.096 | 0/16 |
| TNBC FBI | SA530 | 17685 | 324 | 1 | 9408 | 3 | 0.092 | 0/9 |
| | SA609 | 44170 | 5993 | 24 | 37237 | 7 | 0.019 | 15/49 |
| TNBC HRD-Dup | SA501 | 23098 | 2473 | 36 | 18399 | 3 | 0.039 | 3/9 |
| | SA535 | 25218 | 1801 | 2 | 17875 | 8 | 0.040 | 6/16 |

***Table S1*** Summary of meta-cohort of 4 triple negative breast cancer and 8 high-grade serous ovarian cancer samples from Funnell *et al.* [19].

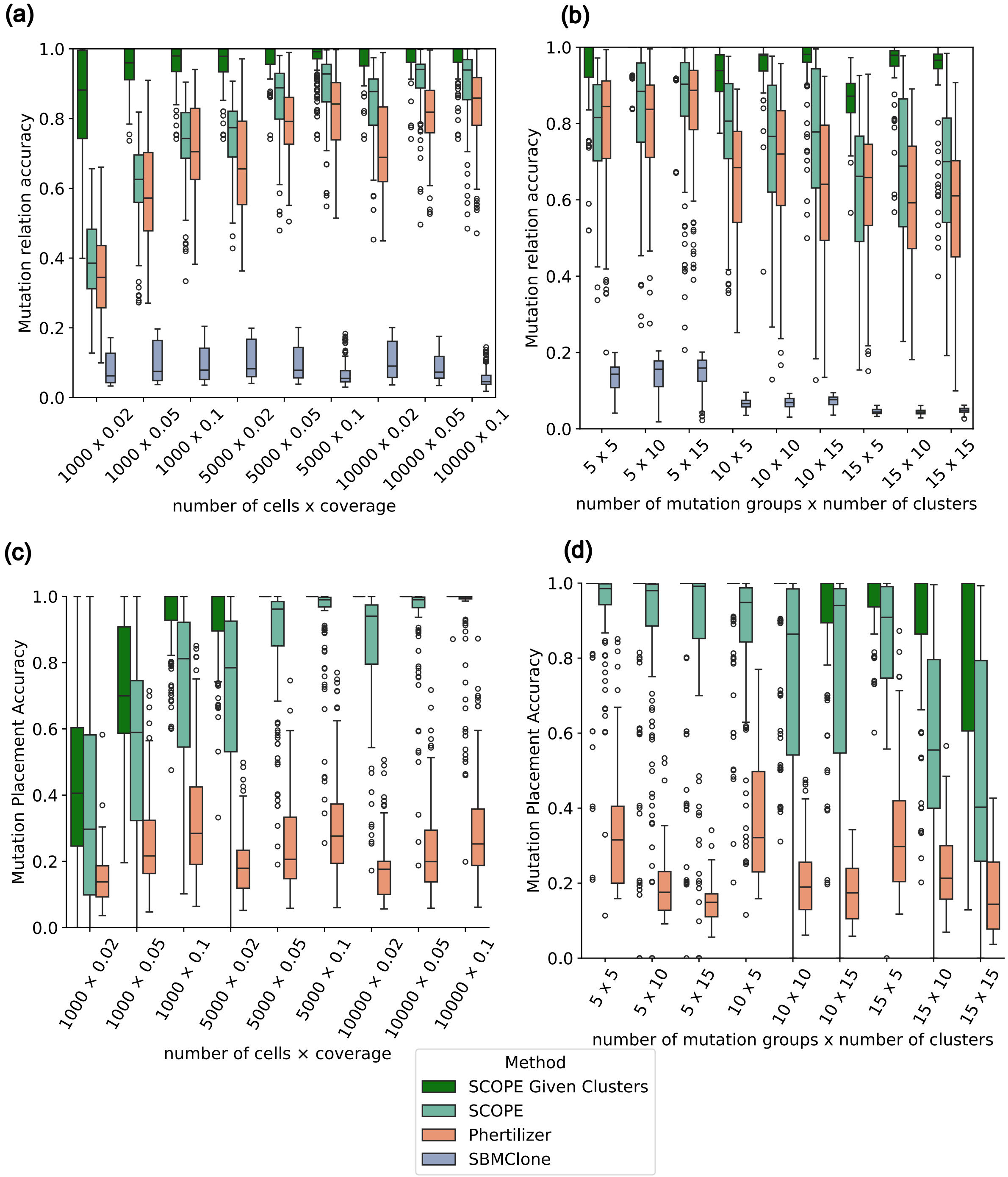

***Fig. S3*** **SCOPE outperforms existing methods on simulated data**. Performance of SCOPE with ground-truth mutation groups, SCOPE (unsupervised), Phertilizer and SBMClone for different setting of simulation parameters. (a) Mutation relationship accuracy for varying number of cell and coverage. (b) Mutation relationship accuracy for varying number of mutation groups and copy-number clusters. (c) Mutation placement accuracy for varying number of cell and coverage. (d) Mutation placement accuracy for varying number of mutation groups and copy-number clusters.
